# Transcriptomic analysis of sorted lung cells revealed a proviral activity of the NF-κB pathway towards SARS-CoV-2

**DOI:** 10.1101/2022.02.25.481978

**Authors:** Anvita Bhargava, Ugo Szachnowski, Maxime Chazal, Dominika Foretek, Sophie-Marie Aicher, Juliana Pipoli da Fonseca, Patricia Jeannin, Guillaume Beauclair, Marc Monot, Antonin Morillon, Nolwenn Jouvenet

## Abstract

Investigations of cellular responses to viral infection are commonly performed on mixed populations of infected and uninfected cells or using single-cell RNA sequencing, leading to inaccurate and low-resolution gene expression interpretations. Here, we performed deep polyA^+^ transcriptome analyses and novel RNA profiling of SARS-CoV-2 infected lung epithelial cells, sorted based on the expression of the viral spike (S) protein. Infection caused a massive reduction in mRNAs and lncRNAs, including transcripts coding for antiviral factors, such as interferons (IFN). This absence of IFN signaling probably explained the poor transcriptomic response of bystander cells co-cultured with S^+^ ones. NF-κB pathway and the inflammatory response escaped the global shutoff in S^+^ cells. Functional investigations revealed the proviral function of the NF-κB pathway and the antiviral activity of CYLD, a negative regulator of the pathway. Thus, our transcriptomic analysis on sorted cells revealed additional genes that modulate SARS-CoV-2 replication in lung cells.

## Introduction

Severe acute respiratory syndrome coronavirus 2 (SARS-CoV-2), which emerged in Wuhan, China, at the end of 2019 is the causative agent of Coronavirus Disease-2019 (COVID-19). Infection may be asymptomatic or it may cause a wide spectrum of symptoms, from mild upper respiratory tract infection to life-threatening pneumonia [1]. Viral replication is not limited to the respiratory tract, but rather occurs in numerous organs, including the blood, heart, vessels, intestines, brain and kidneys [2]. Severity of the disease correlates with an excessive pro-inflammatory immune response [3–5], which may be responsible for the symptoms observed in patients. Inflammation is a vital defense mechanism that is required to initiate an adaptive immune response *via* the recruitment and activation of immune cells. However, the non-resolution of acute inflammation leads to tissue damage [6].

SARS-CoV-2 infection is also characterized by a suppression of interferon (IFN) response in infected cells [7]. IFNs are potent antiviral cytokines secreted by various cell types. The IFN response is initiated by the recognition of viral nucleic acids by cellular receptors. Once activated, these receptors recruit adaptor proteins and kinases that trigger the nuclear translocation of the transcription factors IRF3 and NF-κB, which, in turn, induce the rapid expression of IFNs and proinflammatory cytokines [8]. In particular, type I (IFNα and β) and type III (IFN-λ1 and IFN-λ2/3) IFNs play crucial roles in protecting infected and neighboring cells from virus replication and spread. Once secreted, they will signal in a paracrine and autocrine manner through their receptors, activating a signaling cascade that induces the expression of up to 2000 IFN-stimulated genes (ISGs). Their concerted actions establish the antiviral state by targeting specific stages of viral replication [9,10]. SARS-CoV-2 overcomes IFN responses *via* a wide array of mechanisms involving viral proteins [11–13] and a virus-derived microRNA [14,15]. These viral strategies likely contribute to an impaired IFN response in COVID-19 patients [16] and, consequently, high levels of viral replication.

Several transcriptomic analyses of human cells infected with SARS-CoV-2 have been performed to describe the perturbation of cellular pathways induced by infection, using several cellular models, such as human cells derived from lung, bronchial or colorectal tissue [17–19], as well as post-mortem lung samples of COVID-19 patients [17] and bronchoalveolar lavage fluids (BALF) from patients [20]. These genome-wide investigations of host cellular responses to SARS-CoV-2 infection were performed exclusively using bulk RNA-sequencing (RNA-seq) technologies, *i.e.* by analyzing gene perturbations in mixed populations of infected and uninfected cells. Previous studies on Zika virus infected cells have estimated that only 10% of the repressed and about 30% of the induced genes can be identified in a mixed population containing around one third of infected cells [21]. Bulk transcriptome signals are thus partly drawn into noise background, rendering impossible to efficiently and exhaustively portray the full variation of the host transcripts. The perturbation of cellular responses in SARS-CoV-2 infected and bystander cells have also been analyzed using single-cell (sc) RNA-seq methods. Such studies were performed in a variety of cellular models, including COVID-relevant ones, such as human intestinal organoids [22], human tracheal-bronchial epithelial cells [23,24], human lung cell lines [19] and BALF from patients [25]. However, the technical variability, high noise and massive sample size of scRNA-seq data raise challenges in analyzing the total number of differentially expressed genes (DEGs) [26] out of a limited list of only 1000 to 3000 most expressed genes in individual cells. The balance between the number of cells to be sequenced and the sequencing depth to extract the maximum amount of information from the experiment also affects the results [27].

Moreover, most bulk and single-cell transcriptomic studies performed to investigate the cellular response to SARS-CoV-2 focused on the expression of the referenced coding genome, largely ignoring non-coding and unannotated information, mainly represented by long non-coding RNAs (lncRNAs). These RNAs, which are at least 200 nucleotides (nt) in length, are of specific interest since they play fundamental roles in cellular identity, development and disease progression through epigenetic or post-transcriptional regulation of mRNA expression [28]. Combined RNA-seq data from multiple sources reported over 20 000 lncRNA loci in the human genome [29]. Future studies will plausibly increase this number, since lncRNAs are more cell-type specific [30] and expressed at lower levels than mRNAs [29]. Most of them are independently transcribed by RNA polymerase II and, like protein-coding RNAs, they can be 5’-capped, polyadenylated, and spliced by the cellular machinery [31]. Increasing evidence suggests the involvement of lncRNAs in virus-host interactions and antiviral immunity [32–34]. Current efforts focus on uncovering unannotated RNAs, which could encompass a variety of RNA biotypes, from rare mRNA isoforms to unannotated intergenic long noncoding RNAs, using reference-based approach with the human gencode annotation [35]. However, so far, none of these strategies have been engaged to dissect virus-cell interactions.

Here, we investigated the coding and non-coding transcriptional landscape of lung cells infected with SARS-CoV-2, sorted according to the expression of the viral protein spike (S). Our deep transcriptome analysis using annotated RNA genes and reference-based RNA profiler uncovered pathways that are directly affected by infection and identified coding and non-coding genes contributing to an optimal SARS-CoV-2 replication.

## Results

### Transcriptional landscapes of SARS-CoV-2-infected and bystander lung cells uncovered a global expression shutoff

To analyze transcriptomic changes in infected and bystander cells, human alveolar basal epithelial carcinoma cells (A549) stably expressing the viral receptor ACE2 (A549-ACE2) were infected with a MOI of 1 for 24 hours, fixed, stained intracellularly using antibodies against S proteins and sorted into S-positive (infected cells, S^+^) and S-negative (bystander, S^−^) populations (Fig. 1A and 1B). Around 15% of A549-ACE2 cells were positive for S protein (Fig. 1B). Cells negative for S protein represent either uninfected cells or cells at an early stage of infection, prior to viral protein production. Mock-infected cells served as negative controls. The experiment was performed twice independently in triplicates. PolyA+ RNAs were isolated from mock-infected, S^+^ and S^−^ cells. Around 85% of the total reads mapped to the viral genome in S^+^ cells, while less than 5% of the total reads aligned with the viral genome in S^−^ cells (Fig. 1C), validating our sorting approach. The large dominance of viral reads over cellular reads illustrates the ability of the virus to hijack the cellular machinery for its replication. A similar proportion of SARS-CoV-2 reads in the RNA pool was previously reported in lung epithelial carcinoma Calu-3 cells infected for 8 hours [18]. These differences in the representation of viral RNA between S^+^ and S^−^ cells altered the robustness of the statistical analysis used to identify DEGs. To overcome this limitation, the samples were depleted of viral RNAs (vRNAs) using a set of oligonucleotide probes covering the entire viral genome (Fig. 1A). Following depletion, viral reads represented between 0,01 and 2,8% of the total reads both in S^+^ and S^−^ cells (Fig. 1C).

**Figure 1.**
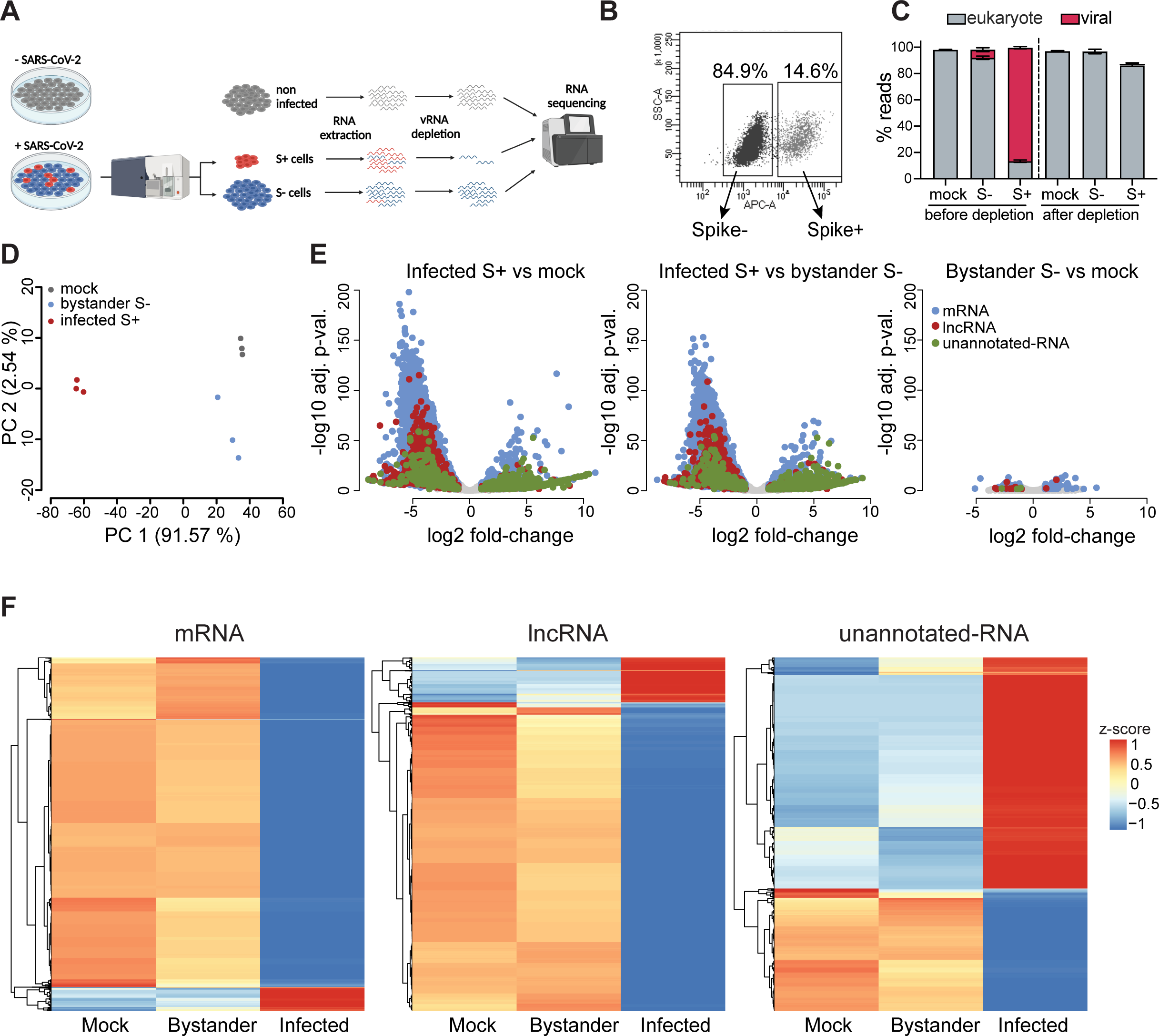
Differential transcriptomic analysis of SARS-CoV-2 infected and bystander lung cells (**A**) Scheme summarizing the experimental workflow. A549-ACE2 cells were infected with SARS-CoV-2 at a MOI of 1 for 24h, stained for viral S protein followed by flow cytometry sorting of productively infected (S+) and bystander (S-) cells. Total RNA from mock, S- and S+ cells was depleted of ribosomal and viral RNAs and sequenced. (**B**) Representative FACS plot of S protein staining used for sorting productively infected cells. (**C**) Percentage of reads in libraries originating from human genome or SARS-CoV-2 sequence, before and after depletion of viral reads. (**D**) PCA plot based on the top 500 most variable genes between mock, bystander (S-) and infected (S+) cells. (**E**) Volcano plots presenting distri-bution of classes of transcripts (mRNA-blue, lncRNA-red, unannotated-green) based on their log2 fold-change for 3 comparisons: infected cells vs mock, infected cells vs bystander and bystander vs mock. (**F**) Heatmaps presenting z-score of log2 normalized counts for all differentially expressed genes between mock, bystander and infected cells, separated for mRNAs, lncRNAs and unannotated RNA.

Coding and long non-coding genes were identified using gencode annotation (v32), while unannotated RNAs were recovered with Scallop assembler [35] (Fig. 1D-F, Fig. S1A-B and tables S1-S3). Principal component analysis (PCA) of polyA+ transcriptomes segregated S^+^ cells from S^−^ and mock-infected ones (Fig. 1D). This segregation based on S expression represented around 92% of the transcriptomic differences between the samples (Fig. 1D). Only subtle differences (2,5%) distinguished bystander and mock-infected cells (Fig. 1D), suggesting that the transcriptional landscapes of these 2 cell populations were very similar. An absence of response of S^−^ cells was unexpected since cytokines, which are commonly secreted by virally infected cells, activate an antiviral state in bystander cells through surface receptors.

Analysis of gene expression allowed identification of thousands of annotated coding and non-coding genes that were differentially expressed (absolute fold change ≥2, p-value < 0.05) in S^+^ cells as compared to S^−^ or mock-infected ones (Fig. 1E-F and tables S1-S3). We identified around 13 times more downregulated coding genes than upregulated ones in S^+^ cells (Fig. 1E-F), suggesting that infection triggers a massive, but incomplete, shutoff of gene expression and/or degradation of host RNAs. Among the top upregulated coding genes in S^+^ cells, we confirmed candidates revealed by previous analyses performed in non-sorted non-vRNA-depleted A549-ACE2 cells, such as CXCL8, CCL20, IL6 and NFKB1 [17,36,37], but also novel highly significant candidates, including IL32 and ITGAM (table S1). The genes encoding IFN type I and type III were not upregulated in S^+^ cells, as compared to mock-infected cells (Fig. S1C). Accordingly, ISGs were not upregulated either in S^+^ cells. This absence of innate immune response in infected cells agrees with previous analyses performed in mixed population of A549-ACE2 cells infected with SARS-CoV-2 [17,36,37]. Such absence of innate response reflects the ability of the virus to potently inhibit the IFN response *via* numerous mechanisms in human cells [12]. Around 1260 annotated lncRNAs were downregulated in S^+^ cells as compared to mock-infected cells, and 184 were upregulated (Fig. 1E-F and table S2). RFPL3S, ADIRF-AS1 and WAKMAR2 were among the top 15 upregulated lncRNAs in S^+^ cells. RFPL3S and ADIRF-AS1 have no known functions, whereas WAKMAR2 restricts NF-kB-induced production of inflammatory chemokines in human keratinocytes [38]. Among the top downregulated lncRNAs, we identified HOXA-AS2 and NKILA, which are negative regulators of NF-kB signaling, in endothelial cells and breast cancer cell lines, respectively [39,40]. Altered expression of WAKMAR2, HOXA-AS2 and NKILA in infected cells could thus play a role in viral-associated inflammation. From the 1400 unannotated transcripts we detected using Scallop assembler [35], around 800 unannotated polyA+ transcripts were also differentially expressed in S+ cells as compared to S^−^ ones (Fig. 1E-F and table S3).

In agreement with the PCA (Fig. 1D), volcano plots and heat maps revealed that S^−^ bystander cells and mock-infected control cells exhibited very similar transcriptomic profiles (Fig. 1E-1F and Fig. S1A-S1B). Only around 170 polyA+ transcripts were differentially expressed in S^−^ cells as compared to mock-infected ones (Fig. 1E and tables S1-S3). As a comparison, over 13000 DEGs were identified in S^+^ as compared to mock-infected cells (Fig. 1E and tables S1-S3). These analyses further suggest that S+ cells present none or very little paracrine signaling response. Among the 69 coding genes that were upregulated in S^−^ cells as compared to mock-infected, 29 were also upregulated in S^+^ cells (Fig. S1B and table S3). Some of these common genes were inflammatory genes, such as IL32, IL6 and CCL20. Among the 39 upregulated coding genes that were unique to S^−^ cells, 16 were ISGs (examples include MX1, APOL1 and IFI6). The expression of these inflammatory genes and ISGs in bystander cells could be induced early in infection, prior to the production of S protein.

Our approach reveals that SARS-CoV-2 infection triggers a major down-regulation of gene expression in A549-ACE2 cells. It also shows that S^−^ cells do not exhibit a strong transcriptional signature despite being cultured with S^+^ cells, suggesting the absence of an efficient paracrine communication.

### Separating lung cells based on the expression of the viral S protein improved discovery of DEGs

To compare our differential deep analyses with known datasets, we analyze publicly available polyA+ RNA-seq raw data of unsorted A549-ACE2 infected with SARS-CoV-2 at a MOI of 0.2 [17]. Viral reads represented around 50% of the total number of reads in these unsorted bulk population of cells [17], which was expectedly less than in A549-ACE2 cells positive for S (Fig. 1C). The 2 analyses shared 150 upregulated protein-coding genes and 238 downregulated ones (Fig. 2A, table S4). The vast majority (about 80%) of the downregulated mRNAs that we identified were classified as ‘unchanged’ in the analysis of unsorted cells (Fig. 2A, table S4). Thus, sorting cells based on S expression and depleting viral RNA allowed the identification of over 30 times more downregulated coding genes than in unsorted cells (Fig. 2A, table S4). The poor sensibility of analysis of mixed cell population in detecting downregulated genes is likely due to the large proportion of non-infected cells, in which the majority of genes remained normally expressed, thus masking any decrease of gene expression in the pool of infected cells. Indeed, an artificial reconstruction of a mixed cell population (80% S^−^ and 20% S^+^) supports this hypothesis (Fig. S2). About 41% of the upregulated protein-coding genes and 16% of downregulated ones that we identified were not detected in conventional RNA-seq analysis of mixed populations [17]. This comparison highlights the accuracy and the depth of our analysis.

**Figure 2.**
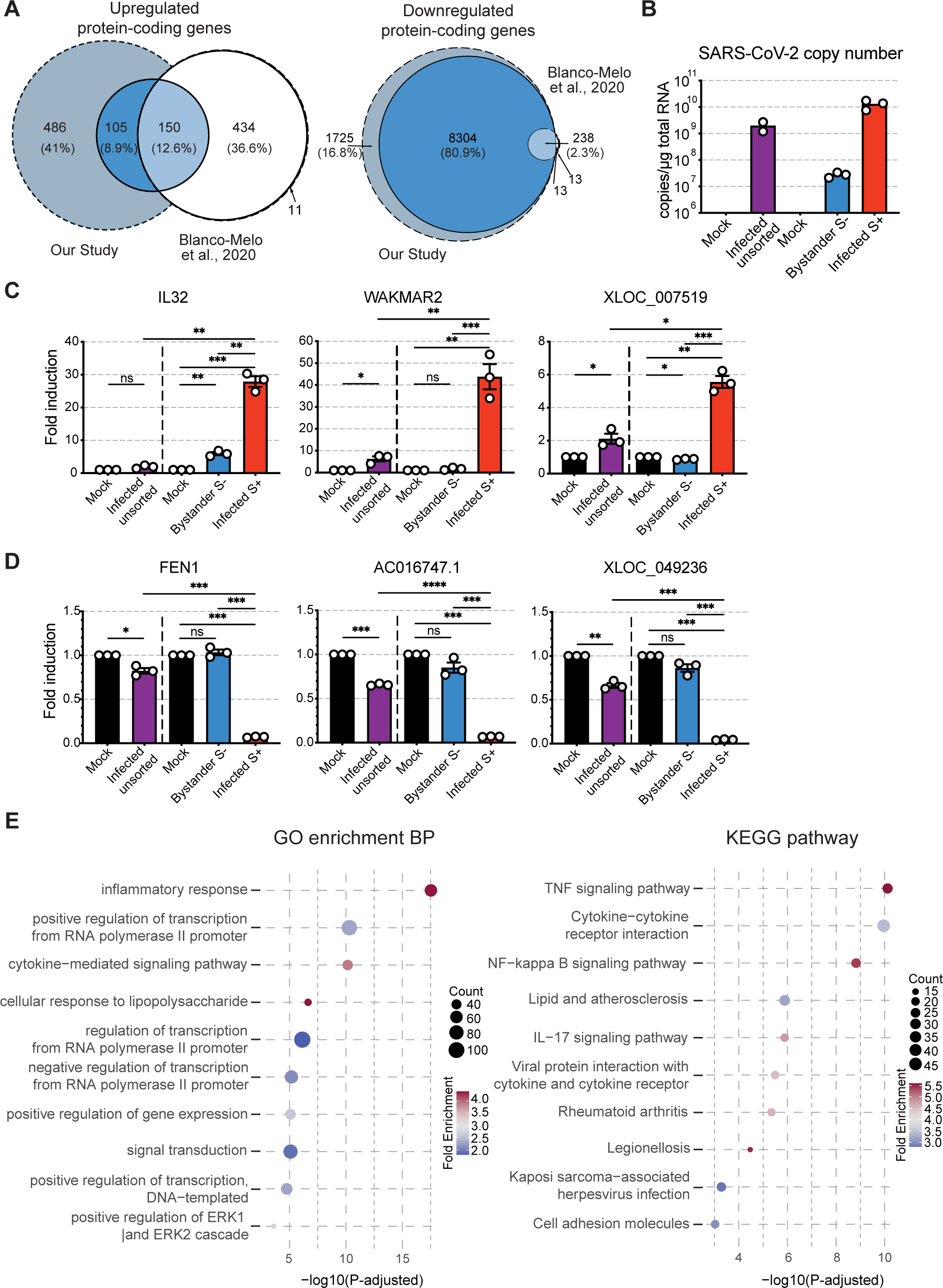
Separating lung cells based on the expression of the viral S protein improved discovery of DEGs. (**A**) Venn diagram repre-senting gene overlap between DE-seq from sorted vs mock samples and mixed vs mock data re-analyzed from Blanco-Melo et al. 2020 (MOI of 0.2). The genes were defined as upregulated if log2 fold change was equal or above 1 (right panel) and equal or below −1 for downregulated genes (left panel). Genes were defined as expressed when they were represented by at least 10 normalized reads in each replicate. The solid lines and central overlap show the genes that appear in both datasets while dashed gray zones outline genes detected in only one of the two datasets. (**B**) RT-qPCR quantification of viral genome copy number per μg of total RNA extrac-ted from A549-ACE2 cells infected by SARS-CoV-2 at a MOI of 1, analyzed either in bulk (left side of graph, n=2 independent expe-riments, line at mean) or post-sorting based on Spike protein expression, allowing distinction between productively infected and bystander subpopulations (right side of graph, n=3 experiments, line at mean). (**C-D**) RT-qPCR quantification of mRNA, lncRNA, and unannotated-RNA, that were identified as upregulated (**C**) or downregulated (**D**) upon infection by SARS-CoV-2 in the RNA-seq analysis, in total RNA extracted from A549-ACE2 cells infected with SARS-CoV-2 at a MOI of 1, analyzed either in bulk (left side of graph) or post-sorting based on Spike protein expression (right side of graph, normalized fold change over mock-infec-ted, n=3 independent experiments, ratio-paired t test, line at mean ± SEM). (**E**) Top 10 enriched GO terms for Biological Process (BP) and KEGG pathways from DAVID database ranked by the adjusted p-value (Benjamini), for upregulated mRNAs identified in RNA-Seq comparison between infected vs mock cells.

To validate the sorting approach combined with vRNA-depletion, we compared mRNA abundances of a few DEGs in a bulk population of cells infected with SARS-CoV-2 for 24 hours, as well as in sorted S^+^ and bystander S^−^ cells infected in the same condition. As expected, S+ cells produced approximately 200-fold more intracellular viral RNAs than did S^−^ cells (Fig. 2B). These qPCR analyses confirm that some S^−^ cells are at an early stage of viral replication, prior to viral protein expression (Fig. 1B). We included in the analysis three coding transcripts (IL32, ITGAM and TRAF1), two lncRNAs (WAKMAR2 and AL132990.1) and one unannotated transcript (XLOC_007519) that were identified amongst the most upregulated RNAs in S^+^ A549-ACE2 cells (Fig. 2C, S3A, S3B and tables S1-S3). The abundance of IL32 mRNA did not increase significantly in the infected bulk population, as compared to mock-infected cells (Fig. 2C). By contrast, IL32 mRNA levels increased around 30-fold in S^+^ cells, compared to those in mock-infected cells (Fig. 2C). This difference explains why IL32 was not identified as an up-regulated gene in previous RNA-seq analysis performed in mixed population of infected A549-ACE2 cells [17,36]. Similarly, the expression of ITGAM, TRAF1, WAKMAR2, AL132990.1 and XLOC_007519 showed a modest increase in the bulk population and a significant increase in S^+^ cells, as compared to mock-infected cells (Fig. 2C and S3B). The decreased expression of transcripts identified as top downregulated hits in the RNA-seq analysis of S+ cells, such as the coding transcripts FEN1 and SNRPF, the lncRNAs AC016747.1, DANCR and TP53TG1, as well as the unannotated RNA XLOC_049236 (Fig. 2D, S3A and S3C), was significantly more pronounced in S^+^ cells than in the mixed population of cells, when compared to mock-infected cells (Fig. 2D and S3C). Analysis of RNA abundances in sorted cells thus highlighted the increased accuracy of our approach, compared to classical methods, in detecting up- and down-regulated genes.

To identify pathways affected by infection in A549-ACE2 cells, we performed Gene Ontology (GO) terms and KEGG pathway enrichment analysis on the upregulated coding genes in S^+^ cells, as compared to mock-infected cells (Fig. 2E). We observed a significant enrichment in several inflammatory signaling pathways, including TNF and NF-κB signatures, which were previously identified in bulk transcriptomic analysis of infected A549 and Calu-3 cells [36,37] and in scRNA-seq analysis of infected colon and ileum organoids [22]. Members of the superfamily of TNF proteins are multifunctional proinflammatory cytokines. NF-κB plays an important role in promoting inflammation, as well as regulating cell proliferation and survival [41]. Activation of NF-κB is one of the signals transduced by the TNF-superfamily members [42]. These inflammatory signatures are also consistent with those observed in peripheral blood immune cells of severe or critical COVID-19 patients [16].

### Inflammatory cytokines, but not IFNs, are produced and secreted by infected cells

We wondered whether the underwhelming response of the bystander S^−^ cell population could be explained by a defect in paracrine communication between S^+^ and S^−^ cells. Despite being present in high abundance in S^+^ cells as compared to mock-infected cells (table S1 and Fig. 2E), inflammatory cytokine transcripts may not be translated. Indeed, initiation of translation seems to be impaired in SARS-CoV-2 infected cells via two potential mechanisms: acceleration of cytosolic cellular mRNA degradation [18] and blockade of the mRNA entry channel of ribosomes by the viral protein Nsp1 [43–45]. Moreover, viral proteins Nsp8 and Nsp9 disrupt protein secretion in HEK293T cells [43], raising the possibility that cytokines are produced but not secreted by S^+^ cells.

To investigate these possibilities, we selected 5 inflammatory chemokines (IL-6, CXCL1, CCL2, CXCL8/IL-8 and CCL20) whose expression was upregulated in S^+^ cells upon infection (table S1) and quantified their intracellular and secreted levels in lysates and supernatants of A549-ACE2 cells infected for 24 hours (Fig. 3A-B). For comparison, A549-ACE2 cells transfected with the immuno-stimulant poly(I:C) were included in the analysis. In a mixed population of S^+^ and S^−^ cells, mRNAs of these 5 cytokines were significantly more abundant than in mock-infected cells (Fig. S4A), in agreement with the increased levels of mRNAs detected in S^+^ cells by RNA-seq (table S1). Their expression was also induced by poly(I:C) (Fig. S4A). All five cytokines were expressed at detectable levels in cells stimulated by viral infection or poly(I:C) (Fig. 3A), indicating that infection does not hamper the translation of the corresponding mRNAs. As expected, based on their mRNA abundance (Fig. S3A), intracellular levels of IL-6, CCL2 and IL-8 significantly increased upon poly(I:C) stimulation as compared to unstimulated control cells (Fig. 3A). By contrast, despite being induced by poly(I:C) downstream signaling (Fig. S4A), CXCL1 and CCL20 levels were comparable in stimulated and unstimulated cells (Fig. 3A). This could be due to a short protein half-life, protein degradation and/or rapid secretion. Intracellular levels of CXCL1 increased significantly upon infection compared to mock-infected cells (Fig. 3A) while intracellular levels of IL-6, CCL2, IL-8 and CCL20 were similar in both conditions. However, all 5 cytokines were significantly more secreted by infected cells than mock-infected ones (Fig. 3B). Infected cells secreted even more IL-6 and CXCL1 than cells stimulated by poly(I:C) (Fig. 3B). Thus, inflammatory cytokines are expressed and secreted by A549-ACE2 cells infected with SARS-CoV-2, which is in line with the excessive inflammatory response reported in other cellular models [17,19,22,24,37] and characteristic of severe cases of COVID-19 [3–5]. The absence of paracrine communication that was revealed by the RNA-seq analysis of S^−^ cells (Fig. 1) is thus unlikely to be linked to a defect in cytokine expression and secretion in S^+^ cells. It may be explained by the apparent absence of IFN signaling. Indeed, and consistently with prior RNA-seq studies conducted in bulk A549-ACE2 cells [17,36,37], we failed to observe a significant IFN-signature in S^+^ cells (table S1, Fig. 2E and Fig. S1D), despite a robust induction of NF-κB activity (Fig. 2E).

**Figure 3.**
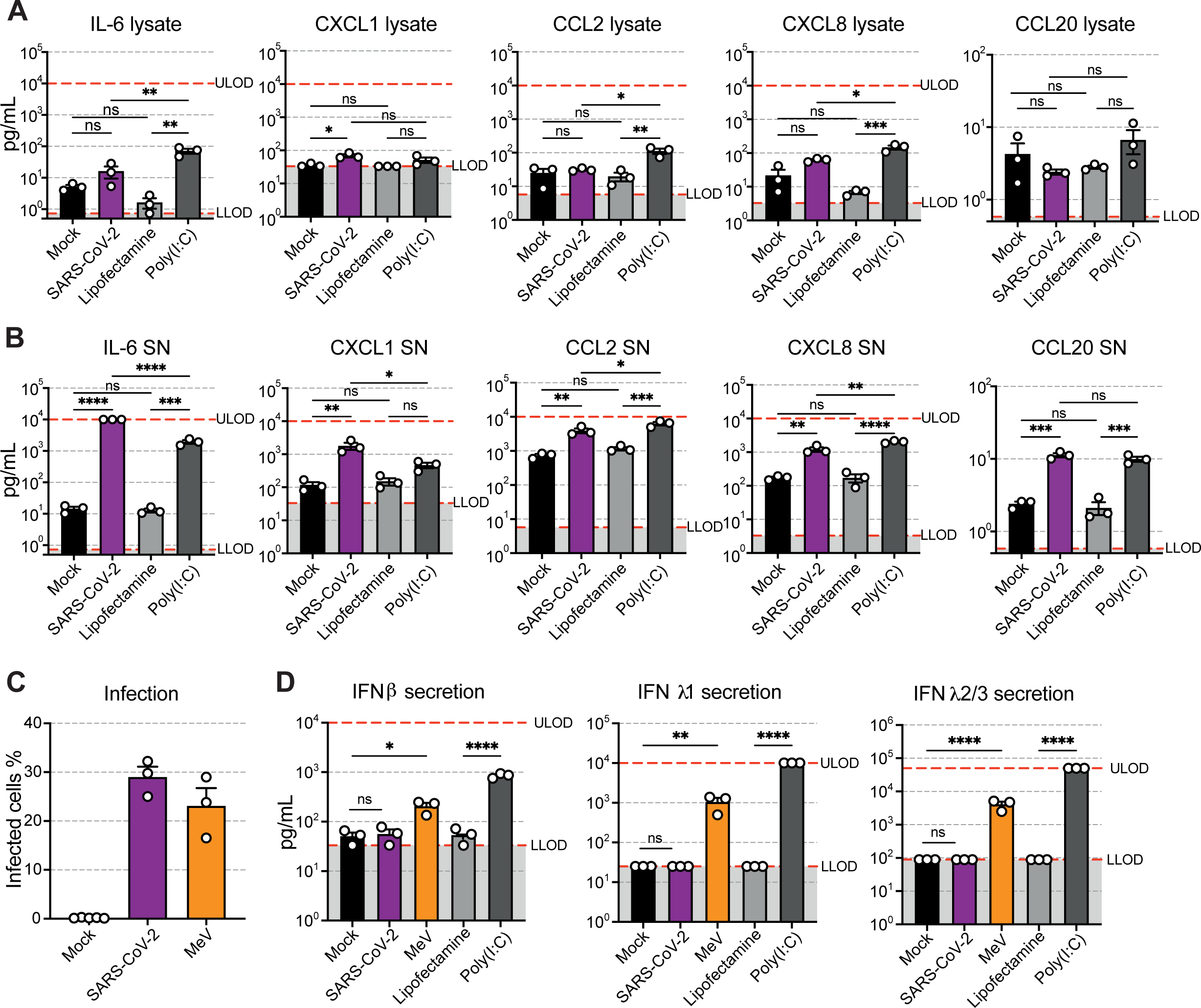
Inflammatory cytokines, but not IFNs, are produced and secreted by infected cells. (**A**) Quantification of the indicated chemokines by cytometry bead array in A549-ACE2 cell lysates obtained 24 hours post mock-infection or infected with SARS-CoV-2 at a MOI of 1, or post-treatment with transfectant alone or in combination with 10 ng/μL of Poly(I:C) (n=3 independent experiments, paired One-Way ANOVA with Turkey’s post-test, line at mean ± SEM). (**B**) Quantification of the indicated chemokines by cytometry bead array in supernatant (SN) of cells shown in (A) (n=3 independent experiments, paired One-Way ANOVA with Turkey’s post-test, line at mean ± SEM). (**C**) Percentages of infected A549-ACE2 cells 24 hours post infection (MOI of 1) with SARS-CoV-2 or Measles virus expressing GFP (MeV), quantified by flow cytometry using Spike protein staining and GFP expression, respectively (n=3 independent experiments, line at mean ± SEM). (**D**) Quantification of secretion of IFNβ, IFNλ1 and IFNλ2/3 by cytometry bead arrays in supernatant of A549-ACE2 cells 24 hours post-infection with SARS-CoV-2 or MeV (MOI of 1), or post-treatment with transfectant alone or in combination with 10 ng/μL of Poly(I:C) (n=3 independent experiments, One-Way ANOVA with Šídák’s post-test, line at mean ± SEM). ULOD: Upper Limit of Detection; LLOD: Lower Limit of Detection.

To validate this further, we compared the level of IFNβ, IFN-λ1 and IFN-λ2/3 transcripts in A549-ACE2 cells infected for 24 hours. Cells treated with poly(I:C) were used as positive controls for IFN production. Cells infected with Measles virus (MeV), a respiratory RNA virus known to trigger an IFN response in A549 cells [46], were also included in the analysis for comparison. Flow cytometry analysis identified on average 20 to 30% of cells positive for viral proteins upon SARS-CoV-2 or MeV infection (Fig. 3C). As expected, the level of IFNβ, IFN-λ1 and IFN-λ2/3 transcripts increased in poly(I:C)-treated cells compared to cells exposed to the transfecting reagent lipofectamine only (Fig. S3B). Amounts of IFNβ, IFN-λ1 and IFN-λ2/3 transcripts were several orders of magnitude higher in MeV-infected cells than in SARS-CoV-2 infected cells (Fig. S4B). Consistently with mRNA level analysis (Fig. S4B), around 200 and 850 pg/ml of IFNβ were secreted by MeV-infected cells and poly(I:C)-treated cells, respectively (Fig. 3D). SARS-CoV-2 infected cells secreted as little as 50 pg/ml of IFNβ, which was similar to the quantity secreted by mock-infected cells and lipofectamine-exposed cells, likely representing baseline levels (Fig. 3D). MeV infected cells secreted around 1000 pg/ml of IFN-λ1 and 5000 pg/ml of IFN-λ2/3 while no IFN-λ was detected in the supernatant of SARS-CoV-2 infected cells (Fig. 3D). This baseline level of IFN type-I secretion and absence of IFN type-III release by SARS-CoV-2-infected cells is likely to be responsible for the lack of paracrine signaling revealed by the RNA-seq analysis (Fig. 1). These analyses indicates that inflammatory cytokines, but not IFNs, are produced and secreted by infected cells.

### Coding and non-coding NF-κB targets escaped the virus-induced cellular shutoff

Numerous genes associated with the NF-κB signaling pathway fall into the category of genes that escaped the virus-induced cellular shutoff (Fig. 2E). To determine which of these genes were directly controlled by NF-κB, we cross-compared the upregulated genes with known NF-κB target genes. Among the 68 upregulated NF-κB-targets in S^+^ cells, we identified cytokines such as CXCL8/IL8 and IL32 (Fig. 4A, S5A and table S5). NFKB1, which codes for the p105/p50 subunit of the transcription factor, and is itself a NF-κB-target gene [47,48], also showed a significant transcriptional induction in S^+^cells (Fig. 4A and S5A), as well as in a bulk population of infected cells (Fig. S5B). Such mechanism generates an auto-regulatory feedback loop in the NF-κB response [47].

**Figure 4.**
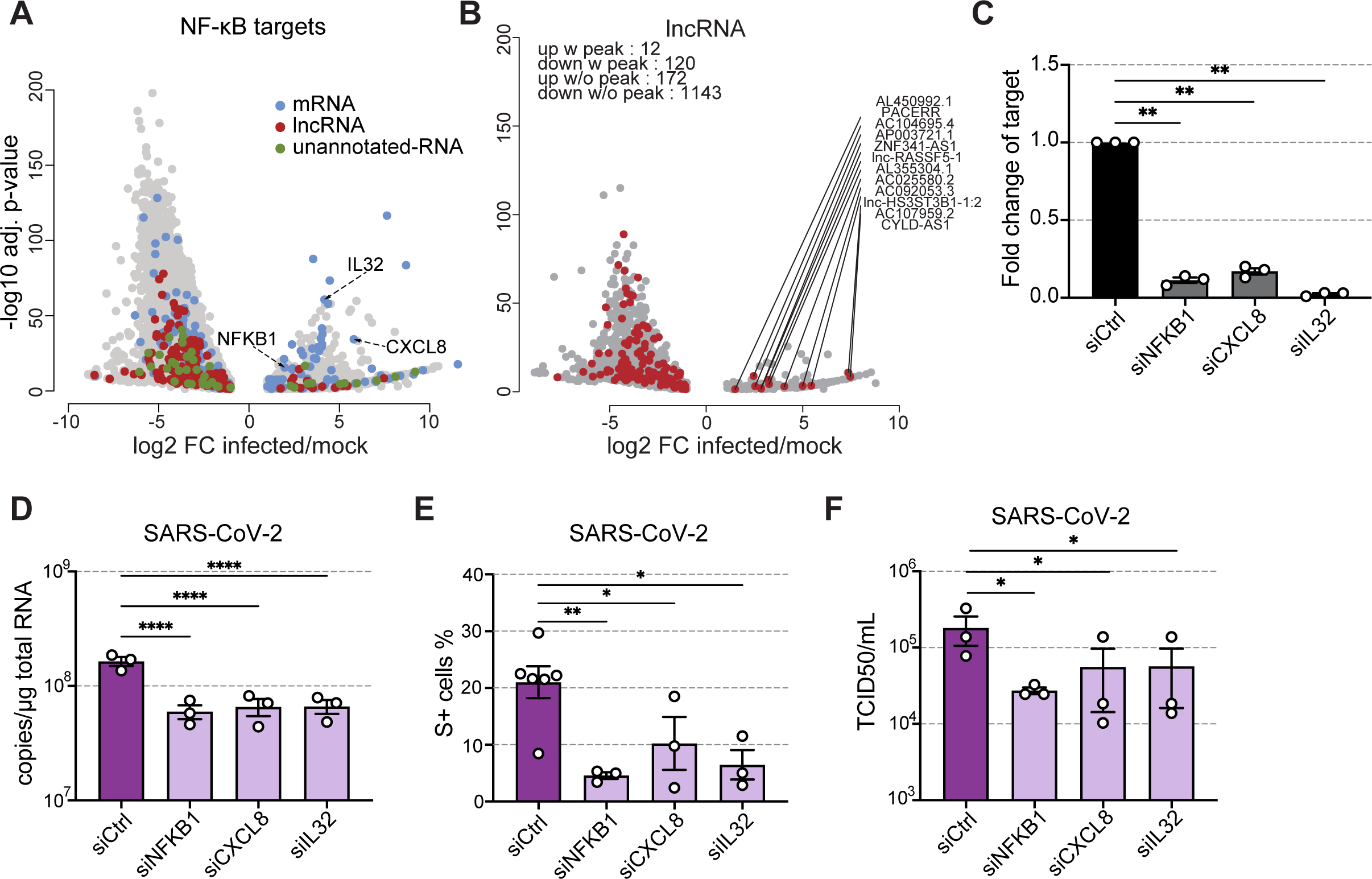
Upregulated NF-κB target genes contribute to an optimal SARS-CoV-2 replication. (**A**) Volcano plot presenting log2 fold change of RNA expression from RNA-seq analysis between S+ and mock cells and showing known NF-κB target mRNAs (labeled in blue), as well as NF-κB target lncRNAs (red) and unannotated RNAs (green) predicted from ChIP and motif analysis. (**B**) Volcano plot presenting log2 fold change of RNA expression from RNA-seq analysis between S+ and mock cells and showing DEGs with peaks of upstream p65 binding, observed via Cut&Run performed after 3 hours TNFα treatment. (**C**) RT-qPCR quantification of knock-down efficiency of indicated transcripts in A549-ACE2 cells, 48 hours post-transfection with a pool of siRNAs targeting indicated genes (normalized fold change over control siRNA, n=3 independent experiments, ratio-paired t test, line at mean ± SEM). (**D**) RT-qPCR quantification of viral genome copy number per μg of total RNA extracted from A549-ACE2 cells, with indicated genes knocked down, 24 hours after infection with SARS-CoV-2 (MOI of 1) (n=3 independent experiments, One-Way ANOVA with Dunnett’s post-test, line at mean ± SEM). (**E**) Percentages of infected A549-ACE2 cells, with selected genes knocked-down, 24 hours post infection with SARS-CoV-2 (MOI of 1), quantified by flow cytometry using Spike protein staining (n=3 independent experiments, mixed model one-Way ANOVA with Dunnett’s post-test, line at mean ± SEM). (**F**) Viral titers in cell supernatants of A549-ACE2 cells, with indicated genes knocked down, 24 hours after infection with SARS-CoV-2 (MOI of 1) (n=3 independent experiments, One-way ANOVA performed on Log-transformed data, with Dunnett’s post-test, line at mean ± SEM). Titers were determined by calculation of TCID50 using the Spearman-Karber Method.

To identify NF-κB-driven lncRNAs, we analyzed NF-κB chromatin immunoprecipitation (ChIP)-sequencing data generated in A549 cells stimulated with TNFα [49] and searched for known NF-κB binding motifs [103]. The analysis recovered 15 NF-κB-targets among the 184 upregulated lncRNAs in S^+^ cells (Fig 4A and table S5), including PACERR, which is known to modulates the expression of NF-κB-target genes *via* a direct interaction with the p50 subunit in U937 macrophages [50]. Novel NF-κB target genes were also identified among unannotated genes (Fig 4A and table S5).

To determine whether the lncRNAs identified by the *in silico* analysis were indeed NF-κB targets, RT-qPCR analysis were performed to assess the levels of three lncRNAs (CYLD-AS1, lnc-RASSF5 and lnc-HS3ST3B1-1:2) following 3 hours of TNFα treatment, and, as a comparison, after 24 hours of SARS-CoV-2 infection (Fig. S6). IL6, CXCL8 and CCL20 were included as positive controls for NF-κB activation. As expected, both TNFα treatment and SARS-CoV-2 infection significantly increased the levels of IL6, CXCL8 and CCL20 mRNAs, as compared to non-treated cells (Fig. S6A). The transcription of CYLD-AS1, lnc-RASSF5 and lnc-HS3ST3B1-1:2 was also induced in TNFα-treated cells (Fig. S6B), as well as in SARS-CoV-2 infected cells, albeit not significantly for lnc-RASSF5 (Fig. S6B). This suggests that transcription of these three lncRNAs occurs via activation of the NF-κB pathway. To confirm that the NF-κB subunit p65 directly binds the upstream promoter of the lncRNAs of interest, a Cut & Run assay was performed to identify direct p65 targets upon TNFα stimulation in A549-ACE2 cells (Fig. 4B). Sequencing results confirmed the presence of direct p65 docking upstream of transcription starts sites (TSS) of the 15 NF-κB-driven lncRNAs identified *in-silico*, except ADIRF-AS1 (Fig. 4B). Thus, coding and non-coding NF-κB targets escaped the virus-induced cellular shutoff.

### The NF-κB signaling pathway is required for an optimal SARS-CoV-2 replication

For functional analysis, we selected NFKB1, CXCL8/IL8 and IL32, which were among the top upregulated NF-κB target genes identified in S^+^ cells (Fig. 4A). NFKB1 served as a positive control in these experiments since reducing its expression was previously shown to decrease SARS-CoV-2 protein expression in A549-ACE2 cells [36]. These results were initially unexpected since NF-κB commonly acts as antiviral factor [41]. Analysis of mRNA abundances showed a significant transcriptional induction of NFKB1 and CXCL8/IL8 in a bulk population of A549-ACE2 cells infected by SARS-CoV-2 for 24 hours, as compared to mock-infected cells (Fig. S5B), validating the RNA-seq analysis performed on S^+^ cells (Fig. 1). We had previously confirmed that IL32 transcripts were significantly more abundant in S^+^ cells than in mock-infected cells (Fig. 2C). We explored the potential ability of NFKB1, CXCL8 and IL32 to modulate the replication of SARS-CoV-2 using siRNA-mediated knock-down approaches. siRNA pools efficiently reduced the expression of their respective targets in A549-ACE2 cells (Fig. 4C). Reduced expression of NFKB1, CXCL8 and IL32 significantly decreased intracellular viral RNA yield twenty-four hours post-infection (Fig. 4D). The number of cells positive for the viral protein S, as well as infectious viral particle production, were also significantly reduced in cells expressing low levels of NFKB1, CXCL8 and IL32, as compared to cells transfected with control siRNA pools (Fig. 4E-F). These results confirmed the pro-SARS-CoV-2 activity of NFKB1 in A459-ACE2 cells [36] and revealed that CXCL8 and IL32 also exhibited significant proviral functions. Thus, our sorting approaches identified upregulated NF-κB target genes that contribute to an optimal SARS-CoV-2 replication.

### CYLD reduced SARS-CoV-2 infection by dampening inflammation

CYLD-AS1 is one of the NF-κB dependent lncRNAs induced by SARS-CoV-2 infection that was detected via the *in silico* approach and validated by the Cut & Run analysis (Fig. 5A) It caught our attention since it arises from the strand opposite to the protein-coding gene CYLD, which is also upregulated by SARS-CoV-2 infection (Table S1). CYLD-AS1 and CYLD share the same promoter for bidirectional transcription, within which we detected a peak for direct p65 binding (Fig. 5A). CYLD has deubiquitinating activity and negatively regulates the NF-κB pathway, thus limiting persistent inflammation [51]. It does so by removing poly-ubiquitin tails from upstream signaling factors such as TRAF2 and TRAF6, thus terminating signal transduction [51]. To investigate the role of CYLD in SARS-CoV-2 infection, we used siRNAs to knock-down its expression prior to infection. Knockdown efficiency was confirmed at the transcript and protein levels (Fig. 5B-C). Consistent with our RNA-Seq data, infection increased the abundance of CYLD, leading to a partial rescue of CYLD transcript levels after siRNA treatment (Fig. 5B). The proportion of S^+^ cells was significantly higher in cells expressing reduced levels of CYLD compared to control cells (Fig. 5D). As expected based on its known function [51], depletion of CYLD led to increased expression levels of the mRNAs of the inflammatory cytokines IL-6, CCL2 and IL-8 /CXCL8 upon infection (Fig. 5E). In agreement with this, less IL-6, CCL2 and IL-8 were secreted in unstimulated cells expressing reduced levels of CYLD compared to control cells (Fig. 5F). These results suggest that CYLD acts as an antiviral factor for SARS-CoV-2, most likely by negatively regulating the proviral NF-κB pathway (Fig. 5G).

**Figure 5.**
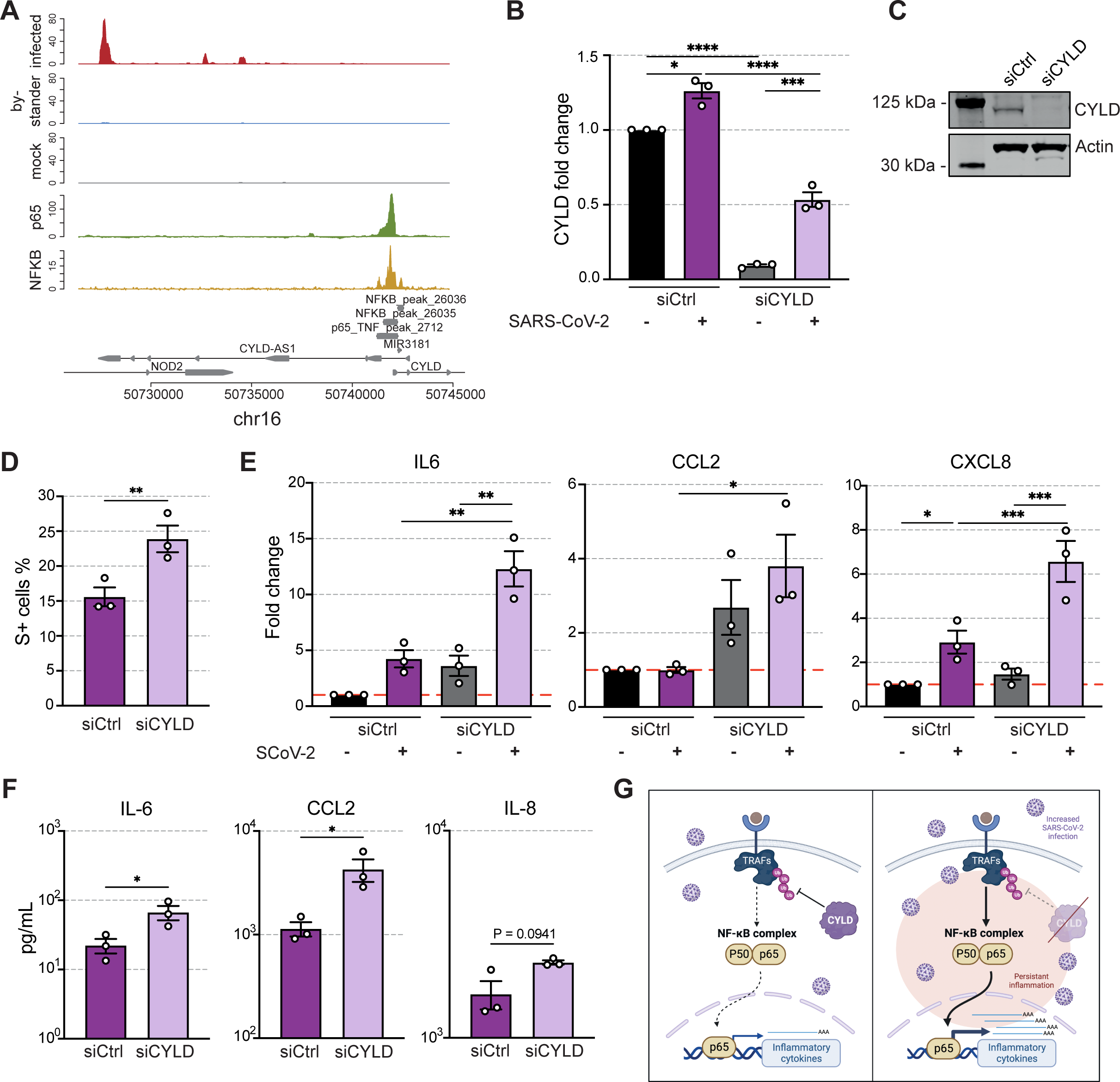
Identification of CYLD gene as a regulator of SARS-CoV-2 infection and inflammation (**A**) Visualization of read coverage (tag/nucleotide) from polyA+ RNA-seq (red, blue, grey), Cut&Run sequencing (green) and ChIP-seq (IP – Input, yellow) at CYLD-AS1 locus. RNA-seq, Cut&Run-seq and ChIP-seq data were normalized independently. (**B**) RT-qPCR quantification of knock-down efficiency of CYLD transcript in A549-ACE2 cells, 72 hours post-transfection with indicated siRNA pools and 24h post-infection with SARS-CoV-2 (MOI of 1) (normalized fold change over control siRNA, n=3 independent experiments, paired One-Way ANOVA with Sidak’s post-test performed on non-normalized ΔCt values, line at mean ± SEM). (**C**) Western Blot showing efficient knockdown of CYLD protein, 48h post-transfection with Ctrl siRNA pool and siRNA pool targeting CYLD transcript. (**D**) Percentages of infected A549-ACE2 cells, with selected genes knocked-down, 24 hours post infection with SARS-CoV-2 (MOI of 1), quantified by flow cytometry using Spike protein staining (n=3 independent experiments, paired t-test, line at mean ± SEM). (**E**) RT-qPCR quantification of transcript induction of indicated genes in A549-ACE2 cells, 72 hours post-transfection with indicated siRNA pools and 24h post-infection with SARS-CoV-2 (MOI of 1) (normalized fold change over mock-infected, control siRNA, n=3 independent experiments, paired One-Way ANOVA with Sidak’s post-test performed on non-normalized ΔCt values, line at mean ± SEM). (**F**) Quantification of secretion of IL-6, CCL-2 and IL-8 by cytometry bead arrays in supernatant of A549-ACE2 cells, 72 hours post-transfection with indicated siRNA pools and 24h post-infection with SARS-CoV-2 (MOI of 1) (n=3 independent experiments, ratio-paired t-test, line at mean ± SEM). (**G**) Working model: CYLD acts by dampening the release of inflammatory cytokines via negative regulation of the NF-κB pathway (left). Reduced expression levels of CYLD (right) lead to increased inflammation and enhanced SARS-CoV-2 infection (Created with BioRender.com).

## Discussion

Transcriptomic analysis of lung A549-ACE2 cells sorted based on Spike expression permitted deep sequencing of many cells synchronized for viral protein expression. Depletion of viral RNA from the samples prior to RNA-seq allowed for a robust identification of host cell DEGs. Our approach thus unveiled an accurate and comprehensive picture of genome-wide signaling networks that are directly affected by SARS-CoV-2 replication in human lung cells. It revealed a massive, but somehow selective, gene expression shutoff in S^+^ cells. Such reduction of cellular transcripts was underestimated in analysis performed on bulk population of infected A549-ACE2 cells [17,36,37] but was detected by RNA-seq analysis performed on bulk population of Calu-3 cells infected at an high MOI [18]. This is probably due to the fact that Calu-3 cells express high levels of ACE2 [52] and are thus naturally permissive to SARS-CoV-2, ensuring a high proportion of infected cells in the mixed culture. SARS-CoV-2 employs several strategies to decrease the level of cellular mRNAs in infected cells, including inhibition of nuclear mRNA export [18,43] and accelerated mRNA degradation as compared to control cells [18]. SARS-CoV-1 and SARS-CoV-2 Nsp1 largely contribute to these processes by interacting with the mRNA export machinery [53] and by inducing endonucleolytic cleavage of the 5′ UTR of capped mRNAs bound to 40S ribosomes [18,54–56]. SARS-CoV-2 RNAs are protected from Nsp1-mediated degradation by their 5’ end leader sequence [18,57], which explains why we observed, in agreement with previous studies performed in A549-ACE2 cells [17] and Calu-3 cells [18], a large dominance of viral RNA over the cellular RNA pool at 24 hpi.

One consequence of this drastic shutoff is the suppression of induction of innate immune genes, such as IFN type I and type III. In agreement with previous RNA-seq studies performed in bulk populations of infected A549-ACE2 cells [17,36] and kidney HEK293T-ACE2 cells [58], our transcriptomic profiling combined with analysis of mRNA levels and IFN secretion showed that infected cells failed to mount an antiviral response. Besides global gene expression reduction in host cells, SARS-CoV-2 has evolved numerous mechanisms to specifically counteract the IFN induction and signaling pathways [12]. For instance, the viral proteins Nsp6 and Nsp13 bind and block the ability of TANK binding kinase 1 (TBK1) to phosphorylate IRF3 [11] and several viral proteins, including the N and Orf6 proteins, dampen STAT1/2 phosphorylation or nuclear translocation [11,59,60]. Consistent with an absence of IFN secretion by S^+^ cells and, consequently, a poor paracrine response, the transcriptome of bystander S^−^ cells largely overlapped with the one of mock-infected cells.

Absence of IFN response is not, however, a universal feature of SARS-CoV-2 infection. Viral replication induces a type I and III IFN response in Calu-3 cells [61–64], primary airway epithelia cultured at the air-liquid interface [61,63], human intestinal epithelial cells [65], organoid-derived bronchioalveolar models [66] and intestinal organoids [67]. When infected at a high MOI, A549-ACE2 cells also induced expression of IFN and ISGs [17]. Thus, *in vitro*, the magnitude of the IFN response elicited by SARS-CoV-2 is cell-type specific and dependent on the viral load. Nevertheless, RNA-seq analysis of postmortem lung tissues from lethal cases of COVID-19 failed to detect IFN-I or IFN-III [17]. Type I IFN responses were highly impaired in peripheral white blood cells of patients with severe or critical COVID-19, as indicated by transcriptional analysis [16]. Moreover, infected patients had no detectable circulating IFN-β, independently of the severity of the disease [16]. Thus, our results corroborate these clinical studies highlighting the efficient shutdown of IFN production by the virus.

Although our RNA-seq analysis identified over 12000 host transcripts that were significantly reduced during SARS-CoV-2 infection as compared to control cells, it also recovered around 1500 transcripts whose levels were significantly elevated and 2800 transcripts whose levels were unchanged upon infection. Among top upregulated genes in S^+^ cells, we identified numerous proinflammatory cytokines, such as IL6, CXCL1, CCL2, IL8/CXCL8 and CCL20. ELISA analysis confirmed that infected cells were producing these inflammatory cytokines. They were previously identified in bulk or sc-RNAseq analysis of A549-ACE2 cells as upregulated [17,36,37], while others, such as IL32, were underreported. High levels of proinflammatory cytokine transcripts have been also observed in infected primary bronchial cells [17], lung macrophages [25] and post-mortem lung samples of COVID-19-positive patients [17]. Thus, SARS-CoV-2 appears to selectively inhibit IFN signaling while allowing chemokine production in lung cells.

GO and KEGG pathway analyses confirmed the upregulation of an inflammatory response in S^+^ cells, including TNF- and NF-κB-transcriptional signatures. An NF-κB transcriptional footprint was previously identified in RNA-seq analysis of bulk population of SARS-CoV-2-infected tracheal-bronchial epithelial cells [24] and in scRNA-seq analysis of infected A549-ACE2 cells [36]. Microarray analysis of Calu-3 cells infected with SARS-CoV-2 also showed a specific bias towards an NF-κB mediated inflammatory response [37]. Finally, inflammatory genes specifically up-regulated in peripheral blood immune cells of severe patients or critical COVID-19 patients mainly belonged to the NF-κB pathway [16]. Consistently, among the 741 upregulated protein-coding genes that we identified in S^+^ cells, 68 possess an NF-κB binding site in their promoter regions. Examples include IL6, CXCL8/IL8 and IL32. We also identified NF-κB binding site in the promoter regions of lncRNAs that were upregulated in S^+^ cells, such as CYLD-AS1 and PACERR, both by *in silico* and Cut-and-Run analysis. NF-κB contribution to the antiviral response is well described and is supported by numerous *in vivo* experiments showing that mice deficient in different NF-κB subunits are more susceptible to viral infection than wild-type mice [41]. Consistently, many viruses have evolved strategies to counteract the NF-κB-mediated antiviral response [68]. However, certain human viruses, such as HIV-1, Epstein-Barr virus and influenza A virus, activate NF-κB to block apoptosis and prolong survival of the host cell to gain time for replication [69]. Our data show that disruption of NF-κB function through silencing of its subunit p105/p50 diminished the production of viral RNAs and proteins at 24 hpi in A549-ACE2 cells, confirming its proviral role [36,37]. Several SARS-CoV-2 proteins could contribute to the activation of NF-κB signaling in infected cells. When individually expressed, Orf7a and Nsp14 activate NF-κB signaling pathway and induce cytokine expression, in Hela and HEK293T cells, respectively [70,71]. Nsp5 also induces the expression of several inflammatory cytokines, such as IL-6 and TNF-α, through activation of NF-κB in Calu-3 and THP1 cells [72]. Further studies are required to understand how SARS-CoV-2 benefits from hijacking NF-κB-driven functions.

Consistent with a proviral role of NF-κB in the context of SARS-CoV-2 infection, we found that diminished expression of two NF-κB target genes (IL32 and CXCL8/IL8,) significantly decreased viral RNA, viral protein production and infectious viral titers. IL-32 is a proinflammatory interleukin secreted by immune and non-immune cells that induces the expression of other inflammatory cytokines, including TNF-*α*, IL-6, and IL1*β* [73]. IL-32 was previously described as an antiviral factor in the context of infection with several RNA and DNA viruses. For instance, its secretory isoform reduces the replication of Hepatitis B virus by stimulating the expression of IFN-λ1 [74]. Its antiviral activity was also demonstrated in U1 macrophages infected with HIV-1 [75] and canine kidney cells infected with influenza A [76], using silencing and over-expression approaches, respectively. Further studies are required to understand the pro-SARS-CoV-2 function of endogenous IL-32. It may support SARS-CoV-2 replication *via* its ability to activate NF-κB [77]. IL-8 is a potent neutrophil chemotactic factor. It was previously shown to possess proviral functions in the context of infection by several unrelated RNA and DNA viruses, probably via inhibition of the antiviral action of IFN-α [78,79]. It could act in a similar manner in SARS-CoV-2 infected A549-ACE2 cells. CYLD, which is an upregulated gene in S+ cells, is a negative regulator of the NF-κB pathway [51] but its role in SARS-CoV-2 has not been documented so far. Our data suggest that it acts as an antiviral factor, further confirming the proviral role of NF-κB in the context of SARS-CoV-2 infection.

As for coding genes, there was a higher proportion of down-*versus* up-regulated lncRNAs in S^+^ cells. GO cannot be extrapolated from lncRNAs since most of them have no known function, indicating the need for future studies in this area. Several RNA-seq and microarray studies have identified hundreds of lncRNAs induced by IFN stimulation or viral infection in diverse human and mice cell types [33,34,80–82]. Analysis of a handful of them has provided a glimpse of the potential regulatory impact of this class of RNAs on the IFN response itself [83] and on ISG expression [33,80,82]. However, the investigation of the precise role of individual lncRNAs in IFN-mediated antiviral response is still in its infancy stage. By analyzing publicly available SARS-CoV-2-infected transcriptome data, several studies recovered lncRNAs that were misregulated upon infection of human lung epithelial cell lines, primary normal human bronchial epithelial cells and BALF [84–87]. However, no lncRNA with a direct action on the life cycle of SARS-CoV-2 has been identified yet.

Finally, our analysis profiled about 600 differentially expressed unannotated polyA+ transcripts in S+ and bystander cells. The identification of these unannotated genes confirms that the human genome is far from being well characterized. Having specific RNAs expressed in particular conditions could open the way for the identification of pro- or anti-viral genes that could be used for better prognosis of at-risk patients or for the follow up of the disease severity.

Our data suggests that the genes that are refractory to the viral-induced shutoff are mostly proviral genes. Understanding the molecular mechanisms underlying the selectivity of the shut-off would be interesting. Since coronavirus Nsp1 induces the cleavage of the 5’UTR of capped transcripts bound to 40S ribosomes, the 5’UTR length and/or structure may affect Nsp1 binding and subsequent degradation. Alternatively, the extent of transcript reduction may be linked to their GC content and/or their lengths, which could affect the specificity of the host RNase that is presumably recruited by Nsp1. Discovering the host RNase responsible for transcript degradation in SARS-CoV-2-infected cells will shed light on the mechanism of selectivity of the viral-induced shutoff.

## Material and Methods

### Cell lines

Human lung epithelial A549-ACE2 cells, which have been modified to stably express ACE2 *via* lentiviral transduction, were generated in the laboratory of Pr. Olivier Schwartz (Institut Pasteur, Paris, France). A549-ACE2 and African green monkey Vero E6 cells (ATCC CRL-1586) were cultured in high-glucose DMEM media (Gibco), supplemented with 10% fetal bovine serum (FBS; Sigma) and 1% penicillin-streptomycin (P/S; Gibco). Cells were maintained at 37°C in a humidified atmosphere with 5% CO_2_.

### Virus and infections

Experiments with SARS-CoV-2 isolates were performed in a BSL-3 laboratory, following safety and security protocols approved by the risk prevention service of Institut Pasteur. The strain BetaCoV/France/IDF0372/2020 was supplied by the National Reference Centre for Respiratory Viruses hosted by Institut Pasteur (Paris, France) and headed by Pr. S. van Der Werf. The human sample from which the strain was isolated has been provided by Dr. X. Lescure and Pr. Y. Yazdanpanah from the Bichat Hospital, Paris, France. Viral stocks were produced by amplification on Vero E6 cells, for 72 h in DMEM 2% FBS. The cleared supernatant was stored at 80°C and titrated on Vero E6 cells by using standard plaque assays to measure plaque-forming units per ml (PFU/ml). A549-ACE2 were infected at MOI of 1 in DMEM without FBS. After 2 h, DMEM with 5% FBS was added to the cells. The Measles Schwarz strain expressing GFP (MeV-GFP) was described previously [88] and was used at an MOI of 1.

### Poly I:C and TNF-α stimulation

Cells were stimulated with 10 ng/µL Poly(I:C) (HMW, #vac-pic Invivogen) using Lipofectamine 3000 Reagent (Thermo Fisher Scientific) according to manufacturer’s protocol. Treatment was maintained for 24 hours, concomitantly with infection. Recombinant human TNF-α (R&D Systems 210-TA/CF) was added directly to cell culture medium at a final concentration of 10 ng/mL and maintained for 3 or 24 hours as indicated.

### Flow cytometry

Cells were detached with trypsin, washed with PBS and fixed in 4% PFA for 30 min at 4°C. Intracellular staining was performed in PBS, 2% BSA, 2mM EDTA and 0.1% Saponin (FACS buffer). Cells were incubated with antibodies recognizing the spike protein of SARS-CoV-2 (anti-S2 H2 162, a kind gift from Dr. Hugo Mouquet, Institut Pasteur, Paris, France) and subsequently with secondary anti-human AlexaFluor-647 antibody (1:1000, A21455 Thermo) for 30 min at 4°C. Data were acquired using Attune NxT Acoustic Focusing Cytometer (Thermo Fisher) and analyzed using FlowJo software.

### Western Blotting

Cells were lysed in radioimmunoprecipitation assay (RIPA) buffer (Sigma-Aldrich) supplemented with a protease and phosphatase inhibitor cocktail (Roche) for 15 minutes at 4°C under agitation. Supernatants were recovered and samples were denatured in 4X Protein Sample Loading Buffer (Li-Cor Bioscience) under reducing conditions (NuPAGE reducing agent, Thermo Fisher Scientific) at 70 °C for 10 minutes. Proteins were separated by SDS-PAGE (NuPAGE 4 to 12% Bis-Tris gel; Invitrogen) and transferred to nitrocellulose membranes (Bio-Rad) using a Trans-Blot Turbo Transfer system (Bio-Rad). Membranes were blocked with PBS-0.1% Tween 20 (PBS-T) containing 5% milk and were incubated with rabbit monoclonal anti-CYLD (1:500, #8462 Cell Signaling) and mouse monoclonal anti-β-actin (1:5,000, A5316; Sigma) diluted in blocking buffer overnight at 4°C. The membranes were then incubated with anti-rabbit/mouse IgG [H+L] DyLight 800/680secondary antibodies diluted in blocking buffer for 1h at room temperature. Images were acquired using an Odyssey CLx infrared imaging system (Li-Cor Bioscience).

### SARS-CoV-2 infected and bystander cell-sorting and RNA extraction on fixed samples for RNA-seq

A549-ACE2 cells were seeded the day prior to infection. Cells were infected with SARS-CoV-2 at MOI 1 or mock infected. Infections were done in two independent repeats with three technical replicates each. At 24 h post infection, cells were detached with trypsin, fixed in 4% PFA for 30 min on ice and stained for spike protein as described above for flow cytometry, with RNasin added to FACS buffer (1:100 dilution) just before use to prevent RNA degradation. Infected cell samples were resuspended in PBS 2%, 25 mM Hepes, 5 mM EDTA (sorting buffer) and sorted at 4°C on a FACSAria Fusion4L Sorter into infected (presence of S protein expression) and bystander (absence of viral protein expression) cell populations. Cells were collected in FBS-coated tubes containing buffer with RNasin to minimize RNA degradation. After sorting, cells were pelleted at 500g for 5 min at 4°C and RNA was extracted with the RecoverAll Total Nucleic Acid Isolation Kit starting at the protease digestion step. Digestion was performed for 15 min at 50°C and 15 min at 80°C in the presence of RNasin. Extraction was performed according to manufacturer’s instructions and the addition of RNAsin to all buffers just before use until final elution of RNA in DNAse-free water. Residual DNA was further digested using DNAse I (Invitrogen AM1906). RNAs were sorted at −80°C until further analysis.

### Library preparation, viral RNA depletion and RNA-sequencing

500-1000 ng of total RNA were depleted of SARS-CoV-2 RNA using custom designed probes. The probes were synthesized using the NC_045512.2 Wuhan-Hu-1 complete genome reference. The design was made by Illumina and is composed of 459 probes, separated into two pools synthetized by IDT. For the SARS-CoV-2 depletion, we mixed both pools and used 1µl of this mix per sample, replacing the Ribozero+ probes at the ribodepletion reaction step of the Illumina Stranded Total RNA prep ligation protocol. The SARS-CoV-2 depleted RNA samples were normalized to 300ng and ERCC Spike was added as recommended by the protocol ERCC RNA Spike-In Control mixes User Guide. The libraries were prepared using the Illumina Stranded mRNA Prep Ligation Reference Guide.

### PolyA+ RNA-sequencing analysis of sorted cells

Dataset consists of 9 paired-end libraries (150 nt), 3 replicates per condition: mock, bystander and infected cells. Adaptors were trimmed with Trim Galore v0.6.4 [89] (wrapper for cutadapt v2.10 [90] and FastQC v0.11.9 [91]), with options --stringency 5 --trim-n -q 20 --length 20 --paired --retain_unpaired. Reads were mapped to a reference containing human genome (hg38), SARS-CoV-2 (NC045512.2) and ERCC sequences. STAR v2.7.3a [92] was used to map the reads, with default parameters. Bam files were then filtered using SAMtools v1.10 [93] to retained reads flagged as primary alignment, and with mapping quality > 30 (option -q 30 -F 0x100 - F 0x800). Read coverage was computed for each strand with bamCoverage (deepTools v3.5.0 [94]) with options --binSize 1 --skipNAs --filterRNAstrand forward/reverse. For the detection of unannotated transcripts, Scallop v0.10.5 [35] was used to reconstruct transcripts, with options --library_type first -- min_transcript_coverage 2 --min_splice_bundary_hits 5 --min_flank_length 5. Scallop was run on each library, and the resulting annotations were merged using cuffmerge v1.0.0 [95], with gencode annotation (v32) as reference (-g option). Then BEDtools v2.29.2 [96] was used to retain only intergenic and antisens transcripts regarding gencode annotation. Gene expression quantification was performed using featureCounts v2.0.0 [97], with options -O -M --fraction -s 2 -p, using a merged annotation of gencode v32, SARS-CoV-2 (NC045512.2), newly annotated transcripts and ERCC transcripts. Subsequent analyses were performed in R v3.6.2 [98]. Differential expression analysis was performed using DESeq2 package [99], after filtering out genes with less than 10 raw counts for all replicates in at least one condition. Gene counts were normalized on ERCC counts, using *estimateSizeFactorsForMatrix* function from DESeq2. All pairwise comparisons were performed (mock vs infected, mock vs bystander and bystander vs infected), and genes were retained as differential if adjusted p-value was < 0.05 and log fold-change > 1 or < −1. All plots were made using custom script, except for heatmaps that were done using pheatmap package (RRID:SCR_016418).

### PolyA+ RNA-sequencing analysis of bulk population of infected cells (from a public dataset)

Fastq files produced in the study of Blanco-Melo et al (2020)[17] were retrieved from GEO repository (GSE147507). Dataset consist of single-end libraries (150 nt). We compared A549-ACE2 “mock” cells (SRR11517680, SRR11517681 & SRR11517682) *versus* A549-ACE2 cells infected with SARS-CoV-2 at MOI 0.2 (SRR11517741, SRR11517742 & SRR11517743). Adaptors were trimmed with Trim Galore v0.6.4 [89], with options --stringency 5 --trim-n -q 20 --length 20. Reads were mapped on a reference containing human genome (hg38) and SARS-CoV-2 (NC045512.2) sequence. Bam files were then filtered using SAMtools v1.10 [93] to retained reads flagged as primary alignment, and with mapping quality > 30 (option -q 30 -F 0x100 -F 0x800). Gene expression quantification was performed using featureCounts v2.0.0 [97], with options -O -M --fraction -s 2, using a merged annotation of gencode v32, SARS-CoV-2 (NC045512.2) and newly annotated transcripts. Gene counts were normalized on the full count matrix, using *estimateSizeFactorsForMatrix* function from DESeq2 [99]. Differential analysis was performed as described above.

### GO enrichment analysis

The GO enrichment and KEGG pathway analysis were performed using DAVID online tool (updated version 2021) [100,101]. Upregulated protein-coding genes from each comparison were taken for the analysis with default background for *Homo sapiens*. GOTERM_BP_DIRECT and KEGG pathway were retained and top 10 results based on adjusted p-value (Benjamini) were plotted using ggplot2 R package (v 3.3.0).

### Identification of NF-κB target genes

A list of coding genes that are known targets of NF-kB is available on Gilmore’s laboratory website (https://www.bu.edu/nf-kb/gene-resources/target-genes). We selected genes from this list that were shown to be direct targets of NF-κB, and for which the gene symbol could be retrieved in gencode annotation (354 genes). For identifying lncRNAs and unreferenced RNAs that possess NF-κB binding site in their promoter, we used p65 ChIP-seq data from GEO dataset GSE34329 [49] - one input file and 2 ChIP replicates, 38nt long reads, single-end. Reads were mapped using bowtie2 v2.4.1 using hg38 as reference, and SAMtools was used to retain the one flagged as primary alignment, with mapping quality > 30, and to remove PCR duplicates (markdup, with -r option). NF-κB binding sites were then detected using macs2 v2.2.7.1 [102], with command callpeak -t ChIP_BamFile1 ChIP_BamFile2 -c input_BamFile -f BAM -g hs -s 38 --keep-dup all. Peaks in the first decile of the −log10(qvalue) value were discarded. NF-kB motif genomic coordinates in the human genome were retrieved using EMBOSS fuzznuc v6.6 [103], using motif 5’-G(3)[AG]N[CT](3)C(2) −3’ [104], on forward and reverse strand (option-complement Y). Peaks and NF-κB motif coordinates were compared using BEDtools [96]; if a motif was contained in a peak, the motif strand was assigned to the peak. LncRNA and un-references transcripts were identified as NF-κB potential targets if their promoter region (1kb before transcript TSS) had a peak containing a motif or a peak for which the −log10(qvalue) was in the top 5%.

### CUT & RUN

The experiment was performed in monoplicate, following the protocol from Henikoff Lab with some adjustments (Skene et al, 2018). Briefly, untreated cells and cells treated for 3h with 10 ng/mL of TNF-α were pelleted after scraping and resuspended in wash buffer (HEPES 20mM, NaCl 150mM, Spermidine 0.5mM) at 10^7^ cells/ml. At room temperature, per sample, 10 μl of Concanavalin A beads slurry washed in bead activation buffer (HEPES 20mM, KCl 10mM, CaCl_2_ 1mM, MnCl_2_ 1mM) was added per 10^6^ cells in the total volume of 1 ml of wash buffer and incubated for 10 min with rotation at 15–20 rpm. Antibodies against H3K27ac (Millipore, MABE647) were diluted 50 times, p65 (Cell Signaling, D14E12) diluted 100 times and IgG (Invitrogen, 31235) 500 times in the buffer (wash buffer with 0.1% digitonin, 2 mM EDTA) and 50 μl of diluted antibody was added to the cells fixed on beads and incubated for 1h. Afterward, the beads were washed twice with 250 µl of digitonin buffer (wash buffer with 0.1% digitonin), resuspended in 52 µl digitonin buffer containing 35ng of pAG-MNase (produced at Institut Curie platform), and incubated 10 min with agitation. Beads were washed twice in 250 µl of ice-cold digitonin buffer, resuspended in 100 µl of ice-cold digitonin buffer, kept on ice for 5 min before adding 2 µl of 100 mM CaCl₂ and incubating for 30 min on ice. The reaction was stopped by addition of 100 µl of stop buffer (NaCl 340mM, EDTA 20mM, EGTA 4mM, RNase A 50 μg/mL, glycogen 50 μg/mL and 0.1% digitonin) and incubated for 10 min at 37°C. Bead suspension was centrifuged at 16k rcf at 4°C for 5 min and beads were recovered for DNA fragments purification with NEB Monarch PCR & DNA purification kit, following the protocol enriching for short DNA fragments. DNA was quantified by Qubit HS DNA kit and samples were sent for library preparation and sequencing to Institut Curie’s NGS platform.

### CUT & RUN data analysis

Adaptors were trimmed with Trim Galore using default parameters for paired-end data. Reads were aligned on human genome (hg38) using Bowtie2 (default parameters) and only properly mapped reads with a mapping quality >=30 were kept. Peaks were detected using macs2, with the option –keep-dup all.

### RNA extraction and RT-qPCR assays

Total RNA was extracted from cells with the NucleoSpin RNA II kit (Macherey-Nagel) according to the manufacturer’s instructions. First-strand complementary DNA (cDNA) synthesis was performed with the RevertAid H Minus M-MuLV Reverse Transcriptase (Thermo Fisher Scientific) using random primers. Quantitative real-time PCR was performed on a real-time PCR system (QuantStudio 6 Flex, Applied Biosystems) with Power SYBR Green RNA-to-CT 1-Step Kit (Thermo Fisher Scientific). Data were analyzed using the 2-ΔΔCT method, with all samples normalized to endogenous BPTF, whose gene expression was confirmed as homogenous across samples by RNA-seq. Genome equivalent concentrations were determined by extrapolation from a standard curve generated from serial dilutions of plasmid encoding a fragment of the RNA-dependent RNA polymerase (RdRp)-IP4 of SARS-CoV-2. Primers used for RT-qPCR analysis are given in table S6.

### siRNA-mediated knockdown

A549-ACE2 cells were transfected using Lipofectamine RNAiMax (Life Technologies) with 10nM of control (#4390843, Ambion) or CXCL8 (L-004756-00, Dharmacon), NFKB1 (L-003520-00, Dharmacon), IL32 (L-015988-00, Dharmacon), ADIRF-AS1 (siTOOLs Biotech) siRNAs following the manufacturer’s instructions. 48h after transfection, cells were infected with SARS-CoV-2 for 24 h.

### Chemokine and Interferon expression and secretion

Cell lysates for intracellular chemokine quantification were obtained *via* repeated freeze-thaw cycles at −80°C of cells suspended in media containing protease inhibitor cocktail (Roche Applied Science) and final centrifugation at 8000g to pellet debris. IL6, CXCL1, CCL2, CXCL8 and CCL20 concentrations in supernatants of from control, infected or stimulated cells, were measured using a custom-designed LEGENDplex Human Panel. Data were acquired on an Attune NxT Flow Cytometer (Thermo Fisher) analyzed with LEGENDplex software (BioLegend). Similarly, IFN-β, IFN-λ1 and IFN-λ2/3 concentrations were measured in undiluted supernatants from control, infected or stimulated cells using a LEGENDplex Human Type 1/2/3 Interferon Panel assay (BioLegend) according to the manufacturer’s protocol.

### TCID50 mediated viral titration

A549-hACE2 cells were infected for 24 hours with MOI 1 of SARS-COV-2. Virus was removed at 3hpi and fresh medium containing 2.5% FBS was added for the remaining time. Supernatants were harvested the following day and stored at −80 °C until assay was performed. Briefly, supernatants were 10-fold serially diluted in DMEM supplemented with 2% FBS. Vero E6 cells and 50-μL potions of serially diluted virus suspensions were deposited in 96-well plate in 8 replicate wells. 5 days later, cells were rinsed, fixed and stained using Crystal Violet (0.2% Crystal Violet in 10% Ethanol and 10% Formaldehyde) at room temperature. CPEs were assessed by calculating the TCID50 values using the Spearman-Karber method [105].

### Statistical analysis

Statistical parameters including the exact value of n, precision measures (as means ± SEM), statistical tests and statistical significance are reported in the figure legends. In figures, asterisks denote statistical significance: *p < 0.05, **p < 0.01, ***p < 0.005, ****p < 0.0001, and “ns” indicates not significant. Statistical analysis was performed in GraphPad Prism 9 (GraphPad Software Inc.).

## Acknowledgments

We thank the French National Reference Centre for Respiratory Viruses hosted by Institut Pasteur (France) and headed by Pr. S. van Der Werf for providing the historical SARS-CoV-2 strain; C. Combredet and F. Tangy (Institut Pasteur) for producing and sharing MeV-GFP; H. Mouquet and C. Planchais (Institut Pasteur) for anti-S antibodies and O. Schwartz (Institut Pasteur) for the A549-ACE2 cells. We are grateful to our team member Felix Streicher for helping design the figure 2A and to all members of our laboratories for helpful discussions. We acknowledge the UTechS Immunology Platform of Institut Pasteur for the use of the cell sorter. The Biomics Platform was supported by France Génomique (ANR-10-INBS-09), IBISA and the Illumina COVID-19 Projects’ offer.

## Supplementary figure legends

**Figure S1.**
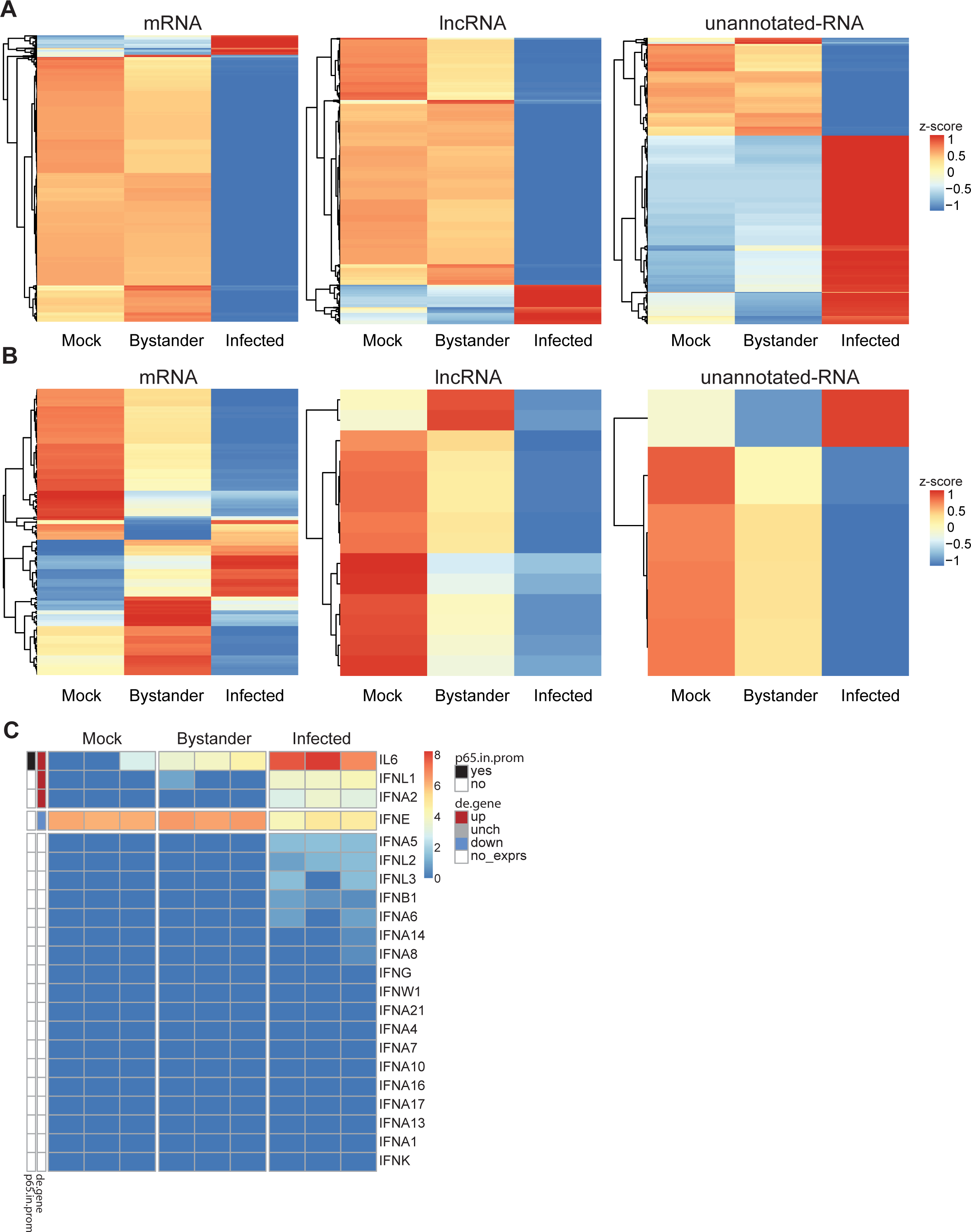
(**A-B**) Heatmaps presenting z-score of log2 normalized counts for differentially expressed genes between (**A**) S+ vs S-cells or (**B**) S-vs mock-infected treated cells, separated for mRNAs, lncRNAs and unannotated RNAs. (**C**) Heatmap showing expression of IFN genes as normalized read counts (log2) of the IFN genes for all replicates in mock, bystander & infected cells. The "de.gene" color scale represents the differentially expressed genes between infected and mock conditions, with "no_exprs" value for genes that were not tested because gene expression was too weak. The "p65.in.prom" color scale indicates whether a gene has a Cut & Run peak (upon TNFα stimulation) in its promoter.

**Figure S2.**
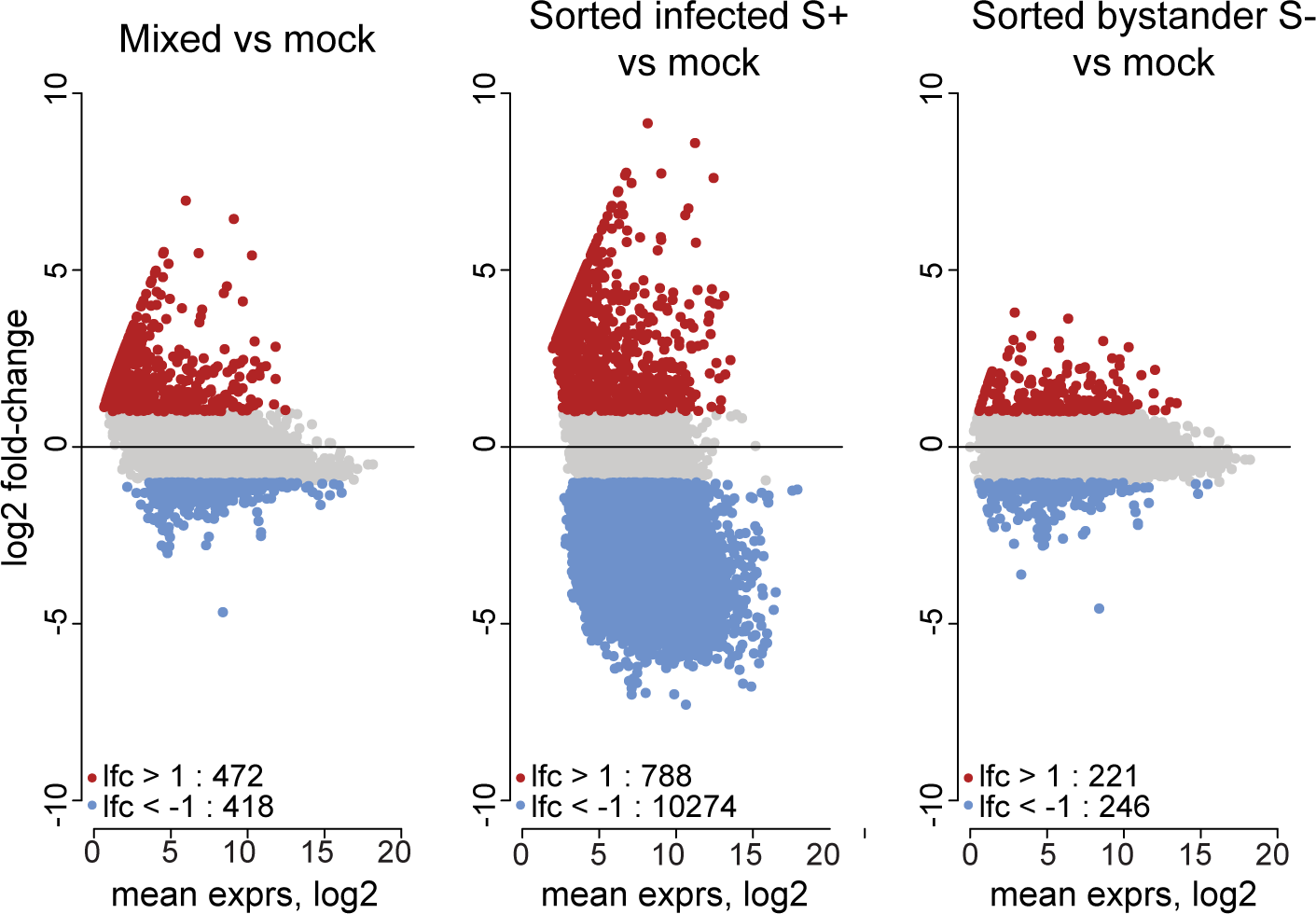
MA plot showing the response to infection of an artificially reconstructed mixed cell population (80% bystander, 20% infected, left) compared to cells sorted based on the expression of the viral protein Spike (infected, middle; bystander, right).

**Figure S3.**
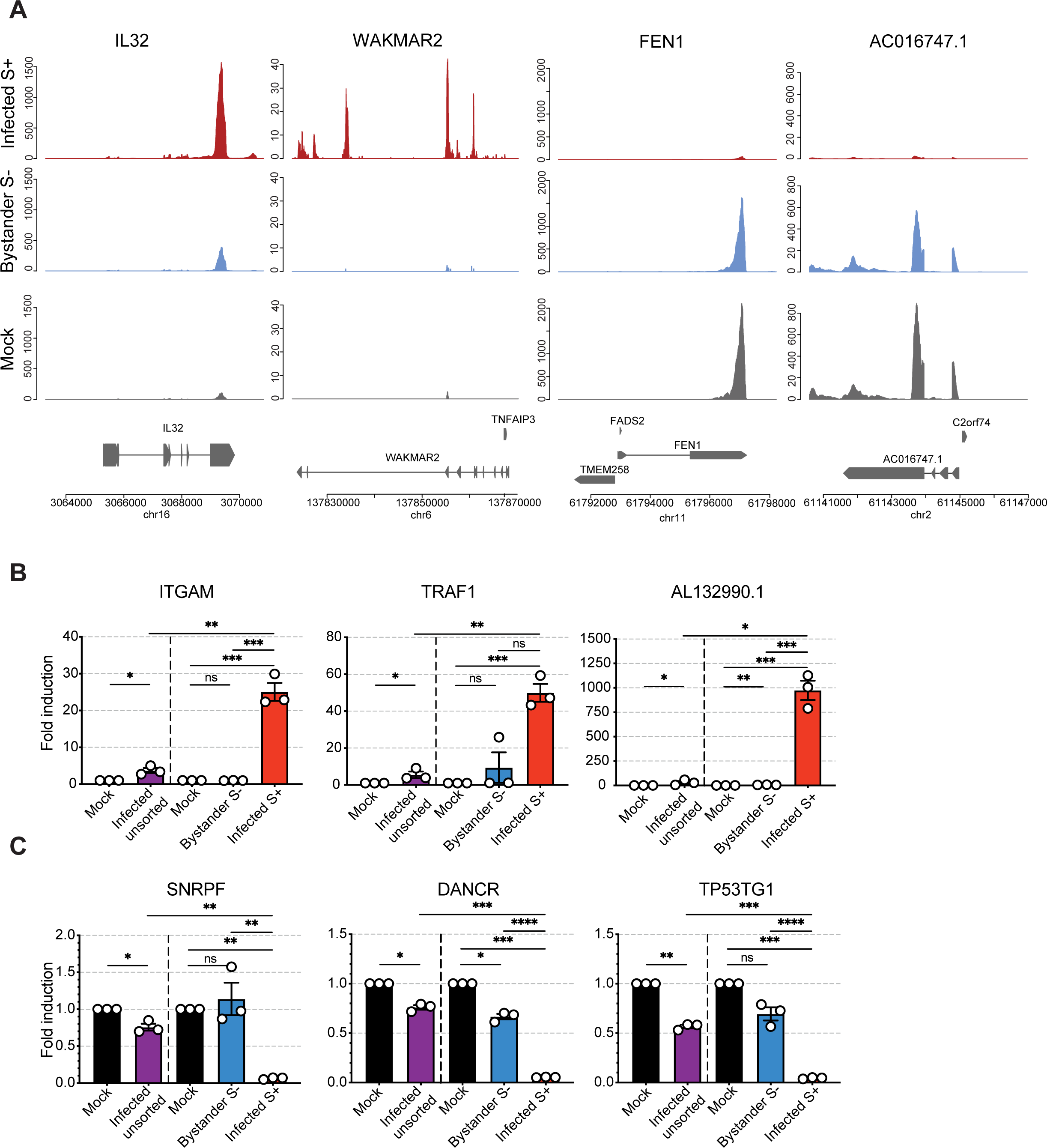
(**A**) Visualization of read coverage (tag/nucleotide) from polyA+ RNA-seq normalized on ERCC reads for IL32, WAKMAR2, FEN1 and AC016747.1. (**B-C**) RT-qPCR quantification of mRNAs and lncRNAs that are either upregulated (**B**) or downregulated (**C**) upon infection with SARS-CoV-2, in total RNA extracted from A549-ACE2 cells infected at an MOI of 1, analyzed either in bulk (left side of graph) or post sorting based on Spike protein (right side of graph, normalized fold change over mock-infected, n=3 independent experiments, ratio-paired t test, line at mean ± SEM).

**Figure S4.**
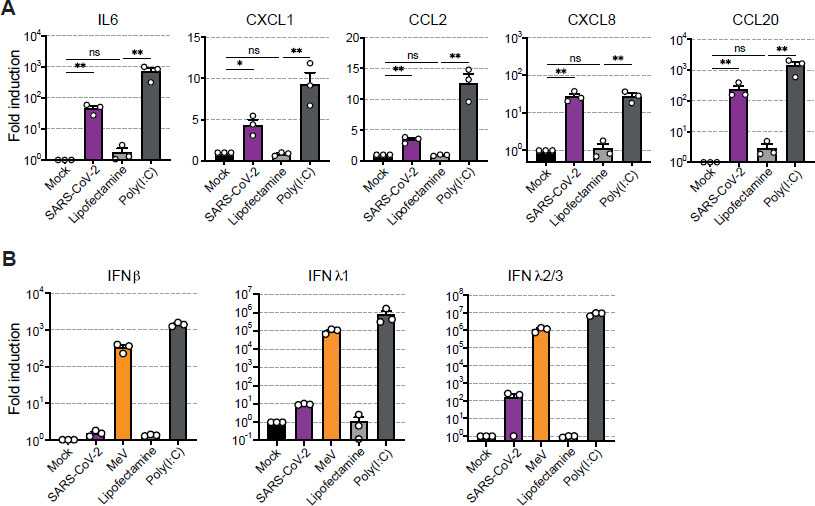
(**A**) RT-qPCR quantification of transcript induction of indicated chemokines in A549-ACE2 cells, 24 hours post infection with SARS-CoV-2 (MOI of 1) or post-treatment with transfectant alone or in combination with 10 ng/μL of Poly(I:C) (normalized fold change over mock-infected, n=3 independent experiments, ratio-paired t test, line at mean ± SEM). (**B**) RT-qPCR quantification of IFNβ, IFNλ1 and IFNλ2/3 transcripts induction in A549-ACE2 cells, 24 hours post infection at an MOI of 1 with SARS-CoV-2 or Measles virus expressing GFP (MeV) or post treatment with transfectant alone or in combination with 10 ng/μL of Poly(I:C) (normalized fold change over mock-infected, n=3 independent experiments, line at mean ± SEM).

**Figure S5.**
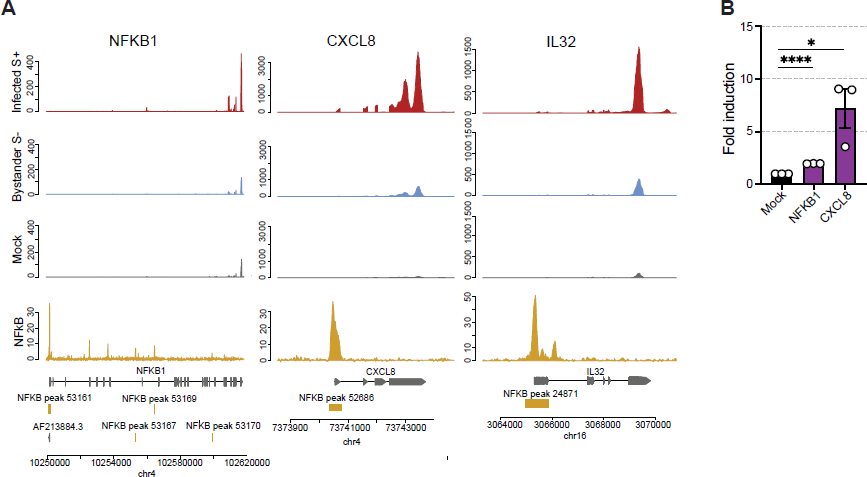
(**A**) Visualization of read coverage (tag/nucleotide) from polyA+ RNA-seq and ChIP-seq (IP – Input) at NFKB1, CXCL8, and IL32 loci. RNA-seq and ChIP-seq data were normalized independently, on ERCC reads for RNA-seq and on library size for ChIP-seq. (**B**) RT-qPCR quantification of NFKB1 and CXCL8 transcripts induction in A549-ACE2 cells, 24 hours post infection with SARS-CoV-2 (MOI of 1) (normalized fold change over mock-infected, n=3 independent experiments, ratio-paired t test, line at mean ± SEM).

**Figure S6.**
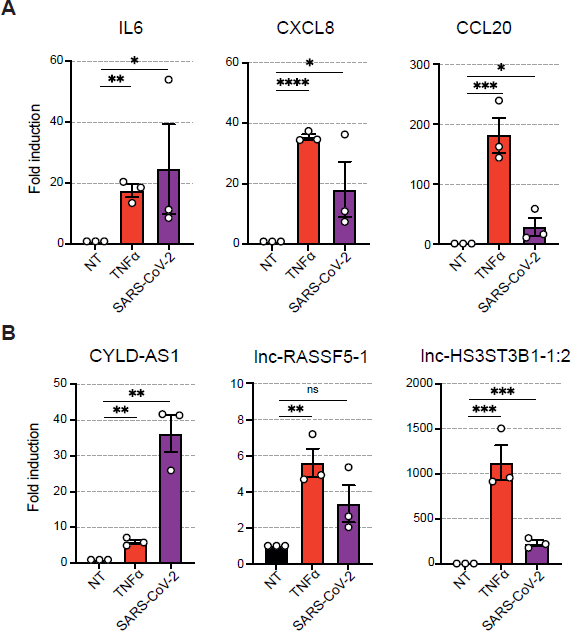
RT-qPCR quantification of three coding transcripts (**A**) and three lncRNAs (**B**) in A549-ACE2 cells, 3 hours post treatment with 10 ng/mL of TNFα or 24 hours post-SARS-CoV-2 infection (MOI of 1) (normalized fold change over mock-infected, n=3 independent experiments, ratio-paired t test, line at mean ± SEM).

## Supplementary tables

Table S1. Differential expression analysis of mRNAs (infected S^+^ vs mock, infected S^+^ vs bystander, and bystander vs mock).

Table S2. Differential expression analysis of lncRNAs (infected S^+^ vs mock, infected S^+^ vs bystander, and bystander vs mock).

Table S3. Differential expression analysis of unannotated RNAs (infected S^+^ vs mock, infected S^+^ vs bystander, and bystander vs mock).

Table S4. Gene overlap between DE-seq from our ‘sorted vs mock’ samples and ‘mixed vs mock’ data re-analyzed from Blanco-Melo *et al.* 2020 (MOI of 0.2).

Table S5. This table shows known NF-κB target mRNAs, as well as predicted NF-κB target lncRNAs and unreferenced RNAs, among upregulated RNAs in S^+^ vs mock cells.

Table S6. RT-qPCR primer sequences

## References

1. Wiersinga WJ, Rhodes A, Cheng AC, Peacock SJ, Prescott HC. Pathophysiology, Transmission, Diagnosis, and Treatment of Coronavirus Disease 2019 (COVID-19): A Review. JAMA. 2020;324: 782–793. doi:10.1001/jama.2020.12839

2. Trypsteen W, Cleemput JV, Snippenberg W van, Gerlo S, Vandekerckhove L. On the whereabouts of SARS-CoV-2 in the human body: A systematic review. PLOS Pathogens. 2020;16: e1009037. doi:10.1371/journal.ppat.1009037

3. Chen G, Wu D, Guo W, Cao Y, Huang D, Wang H, et al. Clinical and immunological features of severe and moderate coronavirus disease 2019. J Clin Invest. 2020;130: 2620–2629. doi:10.1172/JCI137244

4. Lucas C, Wong P, Klein J, Castro TBR, Silva J, Sundaram M, et al. Longitudinal analyses reveal immunological misfiring in severe COVID-19. Nature. 2020;584: 463–469. doi:10.1038/s41586-020-2588-y

5. Giamarellos-Bourboulis EJ, Netea MG, Rovina N, Akinosoglou K, Antoniadou A, Antonakos N, et al. Complex Immune Dysregulation in COVID-19 Patients with Severe Respiratory Failure. Cell Host Microbe. 2020;27: 992-1000.e3. doi:10.1016/j.chom.2020.04.009

6. Chen L, Deng H, Cui H, Fang J, Zuo Z, Deng J, et al. Inflammatory responses and inflammation-associated diseases in organs. Oncotarget. 2017;9: 7204–7218. doi:10.18632/oncotarget.23208

7. Jouvenet N, Goujon C, Banerjee A. Clash of the titans: interferons and SARS-CoV-2. Trends Immunol. 2021;42: 1069–1072. doi:10.1016/j.it.2021.10.009

8. Streicher F, Jouvenet N. Stimulation of Innate Immunity by Host and Viral RNAs. Trends in Immunology. 2019;40: 1134–1148. doi:10.1016/j.it.2019.10.009

9. Schneider WM, Chevillotte MD, Rice CM. Interferon-Stimulated Genes: A Complex Web of Host Defenses. Annu Rev Immunol. 2014;32: 513–545. doi:10.1146/annurev-immunol-032713-120231

10. Schoggins JW. Recent advances in antiviral interferon-stimulated gene biology. F1000Res. 2018;7: 309. doi:10.12688/f1000research.12450.1

11. Xia H, Cao Z, Xie X, Zhang X, Chen JY-C, Wang H, et al. Evasion of Type I Interferon by SARS-CoV-2. Cell Rep. 2020;33: 108234. doi:10.1016/j.celrep.2020.108234

12. Beyer DK, Forero A. Mechanisms of Antiviral Immune Evasion of SARS-CoV-2. J Mol Biol. 2021; 167265. doi:10.1016/j.jmb.2021.167265

13. Sa Ribero M, Jouvenet N, Dreux M, Nisole S. Interplay between SARS-CoV-2 and the type I interferon response. PLoS Pathog. 2020;16: e1008737. doi:10.1371/journal.ppat.1008737

14. Singh M, Chazal M, Quarato P, Bourdon L, Malabat C, Vallet T, et al. A virus-derived microRNA targets immune response genes during SARS-CoV-2 infection. EMBO Rep. 2021; e54341. doi:10.15252/embr.202154341

15. Pawlica P, Yario TA, White S, Wang J, Moss WN, Hui P, et al. SARS-CoV-2 expresses a microRNA-like small RNA able to selectively repress host genes. PNAS. 2021;118. doi:10.1073/pnas.2116668118

16. Hadjadj J, Yatim N, Barnabei L, Corneau A, Boussier J, Smith N, et al. Impaired type I interferon activity and inflammatory responses in severe COVID-19 patients. Science. 2020;369: 718–724. doi:10.1126/science.abc6027

17. Blanco-Melo D, Nilsson-Payant BE, Liu W-C, Uhl S, Hoagland D, Møller R, et al. Imbalanced Host Response to SARS-CoV-2 Drives Development of COVID-19. Cell. 2020;181: 1036-1045.e9. doi:10.1016/j.cell.2020.04.026

18. Finkel Y, Gluck A, Nachshon A, Winkler R, Fisher T, Rozman B, et al. SARS-CoV-2 uses a multipronged strategy to impede host protein synthesis. Nature. 2021;594: 240–245. doi:10.1038/s41586-021-03610-3

19. Wyler E, Mösbauer K, Franke V, Diag A, Gottula LT, Arsiè R, et al. Transcriptomic profiling of SARS-CoV-2 infected human cell lines identifies HSP90 as target for COVID-19 therapy. iScience. 2021;24: 102151. doi:10.1016/j.isci.2021.102151

20. Xiong Y, Liu Y, Cao L, Wang D, Guo M, Jiang A, et al. Transcriptomic characteristics of bronchoalveolar lavage fluid and peripheral blood mononuclear cells in COVID-19 patients. Emerg Microbes Infect. 2020;9: 761–770. doi:10.1080/22221751.2020.1747363

21. Carlin AF, Vizcarra EA, Branche E, Viramontes KM, Suarez-Amaran L, Ley K, et al. Deconvolution of pro- and antiviral genomic responses in Zika virus-infected and bystander macrophages. Proc Natl Acad Sci U S A. 2018;115: E9172–E9181. doi:10.1073/pnas.1807690115

22. Triana S, Metz-Zumaran C, Ramirez C, Kee C, Doldan P, Shahraz M, et al. Single-cell analyses reveal SARS-CoV-2 interference with intrinsic immune response in the human gut. Molecular Systems Biology. 2021;17: e10232. doi:10.15252/msb.202110232

23. Fiege JK, Thiede JM, Nanda HA, Matchett WE, Moore PJ, Montanari NR, et al. Single cell resolution of SARS-CoV-2 tropism, antiviral responses, and susceptibility to therapies in primary human airway epithelium. PLoS Pathog. 2021;17: e1009292. doi:10.1371/journal.ppat.1009292

24. Ravindra NG, Alfajaro MM, Gasque V, Huston NC, Wan H, Szigeti-Buck K, et al. Single-cell longitudinal analysis of SARS-CoV-2 infection in human airway epithelium identifies target cells, alterations in gene expression, and cell state changes. PLOS Biology. 2021;19: e3001143. doi:10.1371/journal.pbio.3001143

25. Liao M, Liu Y, Yuan J, Wen Y, Xu G, Zhao J, et al. Single-cell landscape of bronchoalveolar immune cells in patients with COVID-19. Nat Med. 2020;26: 842–844. doi:10.1038/s41591-020-0901-9

26. Chen G, Ning B, Shi T. Single-Cell RNA-Seq Technologies and Related Computational Data Analysis. Front Genet. 2019;10: 317. doi:10.3389/fgene.2019.00317

27. Zhang MJ, Ntranos V, Tse D. Determining sequencing depth in a single-cell RNA-seq experiment. Nat Commun. 2020;11: 774. doi:10.1038/s41467-020-14482-y

28. Kopp F, Mendell JT. Functional Classification and Experimental Dissection of Long Noncoding RNAs. Cell. 2018;172: 393–407. doi:10.1016/j.cell.2018.01.011

29. Iyer MK, Niknafs YS, Malik R, Singhal U, Sahu A, Hosono Y, et al. The landscape of long noncoding RNAs in the human transcriptome. Nat Genet. 2015;47: 199–208. doi:10.1038/ng.3192

30. Encode Project Consortium. An integrated encyclopedia of DNA elements in the human genome. Nature. 2012;489: 57–74. doi:10.1038/nature11247

31. Guttman M, Amit I, Garber M, French C, Lin MF, Feldser D, et al. Chromatin signature reveals over a thousand highly conserved large non-coding RNAs in mammals. Nature. 2009;458: 223–7. doi:10.1038/nature07672

32. Meng XY, Luo Y, Anwar MN, Sun Y, Gao Y, Zhang H, et al. Long Non-Coding RNAs: Emerging and Versatile Regulators in Host-Virus Interactions. Front Immunol. 2017;8: 1663. doi:10.3389/fimmu.2017.01663

33. Carnero E, Barriocanal M, Segura V, Guruceaga E, Prior C, Borner K, et al. Type I Interferon Regulates the Expression of Long Non-Coding RNAs. Front Immunol. 2014;5:548. doi:10.3389/fimmu.2014.00548

34. Basavappa MG, Ferretti M, Dittmar M, Stoute J, Sullivan MC, Whig K, et al. The lncRNA ALPHA specifically targets chikungunya virus to control infection. Molecular Cell. 2022;82: 3729-3744.e10. doi:10.1016/j.molcel.2022.08.030

35. Shao M, Kingsford C. Accurate assembly of transcripts through phase-preserving graph decomposition. Nat Biotechnol. 2017;35: 1167–1169. doi:10.1038/nbt.4020

36. Nilsson-Payant BE, Uhl S, Grimont A, Doane AS, Cohen P, Patel RS, et al. The NF-κB Transcriptional Footprint Is Essential for SARS-CoV-2 Replication. J Virol. 2021;95: e0125721. doi:10.1128/JVI.01257-21

37. Neufeldt CJ, Cerikan B, Cortese M, Frankish J, Lee J-Y, Plociennikowska A, et al. SARS-CoV-2 infection induces a pro-inflammatory cytokine response through cGAS-STING and NF-κB. Commun Biol. 2022;5: 1–15. doi:10.1038/s42003-021-02983-5

38. Herter EK, Li D, Toma MA, Vij M, Li X, Visscher D, et al. WAKMAR2, a Long Noncoding RNA Downregulated in Human Chronic Wounds, Modulates Keratinocyte Motility and Production of Inflammatory Chemokines. Journal of Investigative Dermatology. 2019;139: 1373–1384. doi:10.1016/j.jid.2018.11.033

39. Zhu X, Liu Y, Yu J, Du J, Guo R, Feng Y, et al. LncRNA HOXA-AS2 represses endothelium inflammation by regulating the activity of NF-κB signaling. Atherosclerosis. 2019;281: 38–46. doi:10.1016/j.atherosclerosis.2018.12.012

40. Liu B, Sun L, Liu Q, Gong C, Yao Y, Lv X, et al. A Cytoplasmic NF-κB Interacting Long Noncoding RNA Blocks IκB Phosphorylation and Suppresses Breast Cancer Metastasis. Cancer Cell. 2015;27: 370–381. doi:10.1016/j.ccell.2015.02.004

41. Santoro MG, Rossi A, Amici C. NF-kappaB and virus infection: who controls whom. EMBO J. 2003;22: 2552–2560. doi:10.1093/emboj/cdg267

42. Aggarwal BB. Signalling pathways of the TNF superfamily: a double-edged sword. Nat Rev Immunol. 2003;3: 745–756. doi:10.1038/nri1184

43. Banerjee AK, Blanco MR, Bruce EA, Honson DD, Chen LM, Chow A, et al. SARS-CoV-2 Disrupts Splicing, Translation, and Protein Trafficking to Suppress Host Defenses. Cell. 2020;183: 1325-1339.e21. doi:10.1016/j.cell.2020.10.004

44. Lapointe CP, Grosely R, Johnson AG, Wang J, Fernández IS, Puglisi JD. Dynamic competition between SARS-CoV-2 NSP1 and mRNA on the human ribosome inhibits translation initiation. Proc Natl Acad Sci U S A. 2021;118: e2017715118. doi:10.1073/pnas.2017715118

45. Yuan S, Peng L, Park JJ, Hu Y, Devarkar SC, Dong MB, et al. Nonstructural Protein 1 of SARS-CoV-2 Is a Potent Pathogenicity Factor Redirecting Host Protein Synthesis Machinery toward Viral RNA. Mol Cell. 2020;80: 1055-1066.e6. doi:10.1016/j.molcel.2020.10.034

46. Helin E, Vainionpää R, Hyypiä T, Julkunen I, Matikainen S. Measles virus activates NF-kappa B and STAT transcription factors and production of IFN-alpha/beta and IL-6 in the human lung epithelial cell line A549. Virology. 2001;290: 1–10. doi:10.1006/viro.2001.1174

47. Oeckinghaus A, Ghosh S. The NF-κB Family of Transcription Factors and Its Regulation. Cold Spring Harb Perspect Biol. 2009;1: a000034. doi:10.1101/cshperspect.a000034

48. Ten RM, Paya CV, Israël N, Le Bail O, Mattei MG, Virelizier JL, et al. The characterization of the promoter of the gene encoding the p50 subunit of NF-kappa B indicates that it participates in its own regulation. EMBO J. 1992;11: 195–203.

49. Raskatov JA, Meier JL, Puckett JW, Yang F, Ramakrishnan P, Dervan PB. Modulation of NF-κB-dependent gene transcription using programmable DNA minor groove binders. Proc Natl Acad Sci U S A. 2012;109: 1023–1028. doi:10.1073/pnas.1118506109

50. Krawczyk M, Emerson BM. p50-associated COX-2 extragenic RNA (PACER) activates COX-2 gene expression by occluding repressive NF-κB complexes. Elife. 2014;3: e01776. doi:10.7554/eLife.01776

51. Kovalenko A, Chable-Bessia C, Cantarella G, Israël A, Wallach D, Courtois G. The tumour suppressor CYLD negatively regulates NF-kappaB signalling by deubiquitination. Nature. 2003;424: 801–805. doi:10.1038/nature01802

52. Murgolo N, Therien AG, Howell B, Klein D, Koeplinger K, Lieberman LA, et al. SARS-CoV-2 tropism, entry, replication, and propagation: Considerations for drug discovery and development. PLOS Pathogens. 2021;17: e1009225. doi:10.1371/journal.ppat.1009225

53. Zhang K, Miorin L, Makio T, Dehghan I, Gao S, Xie Y, et al. Nsp1 protein of SARS-CoV-2 disrupts the mRNA export machinery to inhibit host gene expression. Science Advances. 2021 [cited 6 Feb 2022]. doi:10.1126/sciadv.abe7386

54. Huang C, Lokugamage KG, Rozovics JM, Narayanan K, Semler BL, Makino S. SARS coronavirus nsp1 protein induces template-dependent endonucleolytic cleavage of mRNAs: viral mRNAs are resistant to nsp1-induced RNA cleavage. PLoS Pathog. 2011;7: e1002433. doi:10.1371/journal.ppat.1002433

55. Kamitani W, Huang C, Narayanan K, Lokugamage KG, Makino S. A two-pronged strategy to suppress host protein synthesis by SARS coronavirus Nsp1 protein. Nat Struct Mol Biol. 2009;16: 1134–1140. doi:10.1038/nsmb.1680

56. Kamitani W, Narayanan K, Huang C, Lokugamage K, Ikegami T, Ito N, et al. Severe acute respiratory syndrome coronavirus nsp1 protein suppresses host gene expression by promoting host mRNA degradation. PNAS. 2006;103: 12885–12890. doi:10.1073/pnas.0603144103

57. Mendez AS, Ly M, González-Sánchez AM, Hartenian E, Ingolia NT, Cate JH, et al. The N-terminal domain of SARS-CoV-2 nsp1 plays key roles in suppression of cellular gene expression and preservation of viral gene expression. Cell Reports. 2021;37: 109841. doi:10.1016/j.celrep.2021.109841

58. Sun G, Cui Q, Garcia G, Wang C, Zhang M, Arumugaswami V, et al. Comparative transcriptomic analysis of SARS-CoV-2 infected cell model systems reveals differential innate immune responses. Sci Rep. 2021;11: 17146. doi:10.1038/s41598-021-96462-w

59. Mu J, Fang Y, Yang Q, Shu T, Wang A, Huang M, et al. SARS-CoV-2 N protein antagonizes type I interferon signaling by suppressing phosphorylation and nuclear translocation of STAT1 and STAT2. Cell Discov. 2020;6: 1–4. doi:10.1038/s41421-020-00208-3

60. Miorin L, Kehrer T, Sanchez-Aparicio MT, Zhang K, Cohen P, Patel RS, et al. SARS-CoV-2 Orf6 hijacks Nup98 to block STAT nuclear import and antagonize interferon signaling. Proc Natl Acad Sci U S A. 2020;117: 28344–28354. doi:10.1073/pnas.2016650117

61. Rebendenne A, Valadão ALC, Tauziet M, Maarifi G, Bonaventure B, McKellar J, et al. SARS-CoV-2 triggers an MDA-5-dependent interferon response which is unable to control replication in lung epithelial cells. J Virol. 2021; JVI.02415-20. doi:10.1128/JVI.02415-20

62. Banerjee A, El-Sayes N, Budylowski P, Jacob RA, Richard D, Maan H, et al. Experimental and natural evidence of SARS-CoV-2-infection-induced activation of type I interferon responses. iScience. 2021;24: 102477. doi:10.1016/j.isci.2021.102477

63. Yin X, Riva L, Pu Y, Martin-Sancho L, Kanamune J, Yamamoto Y, et al. MDA5 Governs the Innate Immune Response to SARS-CoV-2 in Lung Epithelial Cells. Cell Rep. 2021;34: 108628. doi:10.1016/j.celrep.2020.108628

64. Lei X, Dong X, Ma R, Wang W, Xiao X, Tian Z, et al. Activation and evasion of type I interferon responses by SARS-CoV-2. Nat Commun. 2020;11: 3810. doi:10.1038/s41467-020-17665-9

65. Stanifer ML, Kee C, Cortese M, Zumaran CM, Triana S, Mukenhirn M, et al. Critical Role of Type III Interferon in Controlling SARS-CoV-2 Infection in Human Intestinal Epithelial Cells. Cell Reports. 2020;32: 107863. doi:10.1016/j.celrep.2020.107863

66. Lamers MM, van der Vaart J, Knoops K, Riesebosch S, Breugem TI, Mykytyn AZ, et al. An organoid-derived bronchioalveolar model for SARS-CoV-2 infection of human alveolar type II-like cells. EMBO J. 2021;40: e105912. doi:10.15252/embj.2020105912

67. Stanifer ML, Kee C, Cortese M, Zumaran CM, Triana S, Mukenhirn M, et al. Critical Role of Type III Interferon in Controlling SARS-CoV-2 Infection in Human Intestinal Epithelial Cells. Cell Rep. 2020;32: 107863. doi:10.1016/j.celrep.2020.107863

68. Deng L, Zeng Q, Wang M, Cheng A, Jia R, Chen S, et al. Suppression of NF-κB Activity: A Viral Immune Evasion Mechanism. Viruses. 2018;10: E409. doi:10.3390/v10080409

69. Neufeldt CJ, Cerikan B, Cortese M, Frankish J, Lee J-Y, Plociennikowska A, et al. SARS-CoV-2 infection induces a pro-inflammatory cytokine response through cGAS-STING and NF-κB. 2020 Jul p. 2020.07.21.212639. doi:10.1101/2020.07.21.212639

70. Su C-M, Wang L, Yoo D. Activation of NF-κB and induction of proinflammatory cytokine expressions mediated by ORF7a protein of SARS-CoV-2. Sci Rep. 2021;11: 13464. doi:10.1038/s41598-021-92941-2

71. Li T, Kenney AD, Liu H, Fiches GN, Zhou D, Biswas A, et al. SARS-CoV-2 Nsp14 activates NF-κB signaling and induces IL-8 upregulation. bioRxiv. 2021; 2021.05.26.445787. doi:10.1101/2021.05.26.445787

72. Li W, Qiao J, You Q, Zong S, Peng Q, Liu Y, et al. SARS-CoV-2 Nsp5 Activates NF-κB Pathway by Upregulating SUMOylation of MAVS. Frontiers in Immunology. 2021;12: 4681. doi:10.3389/fimmu.2021.750969

73. Boreika R, Sitkauskiene B. Interleukin-32 in Pathogenesis of Atopic Diseases: Proinflammatory or Anti-Inflammatory Role? Journal of Interferon & Cytokine Research. 2021;41: 235–243. doi:10.1089/jir.2020.0230

74. Li Y, Xie J, Xu X, Liu L, Wan Y, Liu Y, et al. Inducible Interleukin 32 (IL-32) Exerts Extensive Antiviral Function via Selective Stimulation of Interferon λ1 (IFN-λ1) *. Journal of Biological Chemistry. 2013;288: 20927–20941. doi:10.1074/jbc.M112.440115

75. Nold MF, Nold-Petry CA, Pott GB, Zepp JA, Saavedra MT, Kim S-H, et al. Endogenous IL-32 controls cytokine and HIV-1 production. J Immunol. 2008;181: 557–565. doi:10.4049/jimmunol.181.1.557

76. Li W, Sun W, Liu L, Yang F, Li Y, Chen Y, et al. IL-32: A Host Proinflammatory Factor against Influenza Viral Replication Is Upregulated by Aberrant Epigenetic Modifications during Influenza A Virus Infection. The Journal of Immunology. 2010;185: 5056–5065. doi:10.4049/jimmunol.0902667

77. Kim S-H, Han S-Y, Azam T, Yoon D-Y, Dinarello CA. Interleukin-32: a cytokine and inducer of TNFalpha. Immunity. 2005;22: 131–142. doi:10.1016/j.immuni.2004.12.003

78. Khabar KSA, Al-Zoghaibi F, Al-Ahdal MN, Murayama T, Dhalla M, Mukaida N, et al. The α Chemokine, Interleukin 8, Inhibits the Antiviral Action of Interferon α. J Exp Med. 1997;186: 1077–1085.

79. Pollicino T, Bellinghieri L, Restuccia A, Raffa G, Musolino C, Alibrandi A, et al. Hepatitis B virus (HBV) induces the expression of interleukin-8 that in turn reduces HBV sensitivity to interferon-alpha. Virology. 2013;444: 317–328. doi:10.1016/j.virol.2013.06.028

80. Kambara H, Niazi F, Kostadinova L, Moonka DK, Siegel CT, Post AB, et al. Negative regulation of the interferon response by an interferon-induced long non-coding RNA. Nucleic Acids Res. 2014;42: 10668–80. doi:10.1093/nar/gku713

81. Peng X, Gralinski L, Armour CD, Ferris MT, Thomas MJ, Proll S, et al. Unique signatures of long noncoding RNA expression in response to virus infection and altered innate immune signaling. MBio. 2010;1. doi:10.1128/mBio.00206-10

82. Ouyang J, Zhu X, Chen Y, Wei H, Chen Q, Chi X, et al. NRAV, a long noncoding RNA, modulates antiviral responses through suppression of interferon-stimulated gene transcription. Cell Host Microbe. 2014;16: 616–26. doi:10.1016/j.chom.2014.10.001

83. Du M, Yuan L, Tan X, Huang D, Wang X, Zheng Z, et al. The LPS-inducible lncRNA Mirt2 is a negative regulator of inflammation. Nat Commun. 2017;8: 2049. doi:10.1038/s41467-017-02229-1

84. Meydan C, Madrer N, Soreq H. The Neat Dance of COVID-19: NEAT1, DANCR, and Co-Modulated Cholinergic RNAs Link to Inflammation. Frontiers in Immunology. 2020;11. Available: https://www.frontiersin.org/article/10.3389/fimmu.2020.590870

85. Saha C, Laha S, Chatterjee R, Bhattacharyya NP. Co-Regulation of Protein Coding Genes by Transcription Factor and Long Non-Coding RNA in SARS-CoV-2 Infected Cells: An In Silico Analysis. Noncoding RNA. 2021;7: 74. doi:10.3390/ncrna7040074

86. Vishnubalaji R, Shaath H, Alajez NM. Protein Coding and Long Noncoding RNA (lncRNA) Transcriptional Landscape in SARS-CoV-2 Infected Bronchial Epithelial Cells Highlight a Role for Interferon and Inflammatory Response. Genes (Basel). 2020;11: E760. doi:10.3390/genes11070760

87. Mukherjee S, Banerjee B, Karasik D, Frenkel-Morgenstern M. mRNA-lncRNA Co-Expression Network Analysis Reveals the Role of lncRNAs in Immune Dysfunction during Severe SARS-CoV-2 Infection. Viruses. 2021;13: 402. doi:10.3390/v13030402

88. Combredet C, Labrousse V, Mollet L, Lorin C, Delebecque F, Hurtrel B, et al. A Molecularly Cloned Schwarz Strain of Measles Virus Vaccine Induces Strong Immune Responses in Macaques and Transgenic Mice. Journal of Virology. 2003;77: 11546–11554. doi:10.1128/JVI.77.21.11546-11554.2003

89. Babraham Bioinformatics - Trim Galore! [cited 21 Feb 2022]. Available: https://www.bioinformatics.babraham.ac.uk/projects/trim_galore/

90. Babraham Bioinformatics - FastQC A Quality Control tool for High Throughput Sequence Data. [cited 21 Feb 2022]. Available: https://www.bioinformatics.babraham.ac.uk/projects/fastqc/

91. Martin M. Cutadapt removes adapter sequences from high-throughput sequencing reads. EMBnet.journal. 2011;17: 10–12. doi:http://dx.doi.org/10.14806/ej.17.1.200

92. Dobin A, Davis CA, Schlesinger F, Drenkow J, Zaleski C, Jha S, et al. STAR: ultrafast universal RNA-seq aligner. Bioinformatics. 2013;29: 15–21. doi:10.1093/bioinformatics/bts635

93. Li H, Handsaker B, Wysoker A, Fennell T, Ruan J, Homer N, et al. The Sequence Alignment/Map format and SAMtools. Bioinformatics. 2009;25: 2078–9. doi:10.1093/bioinformatics/btp352

94. Ramírez F, Ryan DP, Grüning B, Bhardwaj V, Kilpert F, Richter AS, et al. deepTools2: a next generation web server for deep-sequencing data analysis. Nucleic Acids Res. 2016;44: W160-165. doi:10.1093/nar/gkw257

95. Trapnell C, Williams BA, Pertea G, Mortazavi A, Kwan G, van Baren MJ, et al. Transcript assembly and quantification by RNA-Seq reveals unannotated transcripts and isoform switching during cell differentiation. Nat Biotechnol. 2010;28: 511–515. doi:10.1038/nbt.1621

96. Quinlan AR, Hall IM. BEDTools: a flexible suite of utilities for comparing genomic features. Bioinformatics. 2010;26: 841–2. doi:10.1093/bioinformatics/btq033

97. Liao Y, Smyth GK, Shi W. featureCounts: an efficient general purpose program for assigning sequence reads to genomic features. Bioinformatics. 2014;30: 923–930. doi:10.1093/bioinformatics/btt656

98. R Core Team. R: A Language and Environment for Statistical Computing. Vienna, Austria; 2017.

99. Love MI, Huber W, Anders S. Moderated estimation of fold change and dispersion for RNA-seq data with DESeq2. Genome Biol. 2014;15: 550. doi:10.1186/s13059-014-0550-8

100. Huang DW, Sherman BT, Lempicki RA. Systematic and integrative analysis of large gene lists using DAVID bioinformatics resources. Nat Protoc. 2009;4: 44–57. doi:10.1038/nprot.2008.211

101. Huang DW, Sherman BT, Lempicki RA. Bioinformatics enrichment tools: paths toward the comprehensive functional analysis of large gene lists. Nucleic Acids Res. 2009;37: 1–13. doi:10.1093/nar/gkn923

102. Zhang Y, Liu T, Meyer CA, Eeckhoute J, Johnson DS, Bernstein BE, et al. Model-based Analysis of ChIP-Seq (MACS). Genome Biology. 2008;9: R137. doi:10.1186/gb-2008-9-9-r137

103. EMBOSS: fuzznuc. [cited 21 Feb 2022]. Available: http://emboss.toulouse.inra.fr/cgi-bin/emboss/fuzznuc

104. Chen FE, Huang DB, Chen YQ, Ghosh G. Crystal structure of p50/p65 heterodimer of transcription factor NF-kappaB bound to DNA. Nature. 1998;391: 410–413. doi:10.1038/34956

105. Aicher S-M, Streicher F, Chazal M, Planas D, Luo D, Buchrieser J, et al. Species-Specific Molecular Barriers to SARS-CoV-2 Replication in Bat Cells. J Virol. 2022;96: e0060822. doi:10.1128/jvi.00608-22

