## Supplemental Table 6 for "Transcriptomic analysis of sorted lung cells revealed a proviral activity of the NF-κB pathway towards SARS-CoV-2"

**Table S6**

| Target Gene | Forward 5' - 3' | Reverse 5' - 3' |
| --- | --- | --- |
| BPTF | GGAGAGATGTTGGTCCTTATGGC | CTTTCCTCTGAGGTGTAGGCGT |
| SARS-CoV-2 nsp12 polymerase | GGTAACTGGTATGATTTC | CTGGTCAAGGTTAATATAGG |
| IL32 | TCAAAGAGGGCTACCTGGAGAC | TCTGTTGCCTCGGCACCGTAAT |
| WAKMAR2 | GCACGTGATGAGCGCGTAG | ATGAGCCAGATAGGGGCAGT |
| XLOC_007519 | GGCTGAATGCACATCACCCTA | TGCTTCATGTCTTCCTTCACG |
| FEN1 | AAGGTCACTAAGCAGCACAATG | GTAGCCGCAGCATAGACTTTG |
| AC016747.1 | TGCCAGTACAACAGCCACAA | CAACCAAATCGGGAAAGCCG |
| XLOC_049236 | TGAGGGGCTTGTATAGTGGGA | ATTTCTCCTTCTCCGCAGGC |
| ITGAM | GGAACGCCATTGTCTGCTTTCG | ATGCTGAGGTCATCCTGGCAGA |
| TRAF1 | TTCACCTCCAATGACGAGCC | AAGCTCTCGGGCTAGGAAGA |
| AL132990.1 | GGGGCTACGGGTTGTCTTTT | AATGATGGGCCACTGAGTCC |
| SNRPF | TGTCTGGACATCTGGGTGAA | TCCCCCACAAAAGATGCTAT |
| DANCR | AGCTCCAGGAGTTCGTCTCT | TGGCTTGTGCCTGTAGTTGT |
| TP53TG1 | GCGAACACTTACACCAGTGC | AGGGTTACTCAGACCTGCCA |
| IL6 | TTCTCCACAAGCGCCTTC | GGGCGGCTACATCTTTGGAA |
| CXCL1 | CTGGCTTAGAACAAAGGGGCT | TAAAGGTAGCCCTTGTTTCCCC |
| CCL2 | CCCAAAGAAGCTGTGATCTTCA | TCTGGGGAAAGCTAGGGGAA |
| CXCL8 | GCGCCAACACAGAAATTATTGTAAA | TGCTTGAAGTTTCACTGGCATC |
| CCL20 | AAGTTGTCTGTGTGCGCAAATCC | CCATTCCAGAAAAGCCACAGTTTT |
| IFNB | AAGCAATTGTCCAGTCCCA | TGCATTACCTGAAGGCCAAG |
| IFNL1 | TCAGCTTGAGTGACTCTTCCAAGG | GCCACATTGGCAGGTTCAAATCTC |
| IFNL2 | TCCAGAACCTTCAGCGTCAG | AGGGCCAAAGATGCCTTAGA |
| NFKB1 | GCTTAGGAGGGAGAGCCCA | GGTATGGGCCATCTGCTGTT |
| CYLD | GGTAATCCGTTGGATCGGTCAG | AGTGCCTCTGAAGGTTCCATCC |
| CYLD-AS1 | GACCACACTGCTGTAGGAAGC | TGTAGATGAGAACGGTGTTGCC |
| lnc-RASSF5-1 | CAATGGGAAAACCAGCACGC | TTTGTGACTCACTCCCAGCA |
| lnc-HS3ST3B1-1:2 | ACAGCCCAGTCAGTAAGTGC | GTCTGTCCAGAAAAAGTTCCTGT |
